# iPSC modeling shows uncompensated mitochondrial mediated oxidative stress underlies early heart failure in hypoplastic left heart syndrome

**DOI:** 10.1101/2021.05.09.443165

**Authors:** Xinxiu Xu, Kang Jin, Abha S. Bais, Wenjuan Zhu, Hisato Yagi, Timothy N Feinstein, Phong Nguyen, Joseph Criscione, Xiaoqin Liu, Gisela Beutner, Kalyani B. Karunakaran, Phillip Adams, Catherine K. Kuo, Dennis Kostka, Gloria S. Pryhuber, Sruti Shiva, Madhavi K. Ganapathiraju, George A. Porter, Jiuann-Huey Ivy Lin, Bruce Aronow, Cecilia W. Lo

**Affiliations:** Department of Developmental Biology, University of Pittsburgh, Pittsburgh, PA, USA; Department of Critical Care Medicine, University of Pittsburgh, Pittsburgh, PA, USA; Department of Pharmacology and Chemical Biology, University of Pittsburgh, Pittsburgh, PA, USA; Vascular Medicine Institute, University of Pittsburgh, Pittsburgh, PA, USA; Department of Biomedical Informatics, University of Pittsburgh, Pittsburgh, PA, USA; Department of Anesthesiology, University of Pittsburgh, Pittsburgh, PA, USA; University of Cincinnati, Cincinnati, United States; Biomedical Informatics, Cincinnati Children’s Hospital Research Foundation, Cincinnati, USA; Centre for Cardiovascular Genomics and Medicine, Faculty of Medicine, The Chinese University of Hong Kong, Hong Kong, China; Department of Biomedical Engineering, University of Rochester, Rochester, NY, USA; Fischell Department of Bioengineering, University of Maryland, College Park, MD; Department of Orthopaedics, University of Maryland School of Medicine, Baltimore, MD; Department of Pediatrics, Pediatrics and Environmental Medicine; Pediatrics, Pharmacology and Physiology, and Aab Cardiovascular Research Institute University of Rochester Medical Center, Rochester, NY, USA; Supercomputer Education and Research Centre, Indian Institute of Science, Bangalore, India

**Keywords:** heart failure, Hypoplastic left heart syndrome (HLHS), congenital heart disease (CHD), induced pluripotent stem cell derived cardiomyocytes (iPSC-CM), permeability transition pore (mPTP), endoplasmic reticulum (ER).

## Abstract

Hypoplastic left heart syndrome (HLHS) is a severe congenital heart defect with 30% mortality from heart failure (HF) in the first year of life, but why only some patients suffer early-HF and its cause remain unknown. Modeling using induced pluripotent stem cell-derived cardiomyocytes (iPSC-CM) showed early-HF patient iPSC-CM have increased apoptosis, redox stress, and failed antioxidant response. This was associated with mitochondrial permeability transition pore (mPTP) opening, mitochondrial hyperfusion and respiration defects. Whereas iPSC-CM from patients without early-HF had hyper-elevated antioxidant response with increased mitochondrial fission and mitophagy. Single cell transcriptomics showed dichotomization by HF outcome, with mitochondrial dysfunction and endoplasmic reticulum (ER) stress associated with early-HF. Importantly, oxidative stress and apoptosis associated with early HF were rescued by sildenafil inhibition of mPTP opening or TUDCA suppression of ER stress. Together these findings demonstrate a new paradigm for modeling clinical outcome in iPSC-CM, demonstrating uncompensated mitochondrial oxidative stress underlies early HF in HLHS.

## Introduction

Congenital heart disease is one of the most common birth defects affecting 0.5% of live births (Feinstein et al., 2012). Hypoplastic left heart syndrome (HLHS) is a severe CHD in which the left ventricle (LV) and aorta are small and nonfunctional. While survival with HLHS is made possible by staged surgical palliation that recruits the RV to become the single pumping chamber (Oster et al., 2013), there remains high morbidity and mortality. The10-year transplant free survival stands at only 39-50%(Driscoll et al., 1992; Garcia et al., 2020; Gentles et al., 1997). However, the greatest risk is in the first year of life with 30% mortality reported (Alsoufi et al., 2016; Cleves et al., 2003; Tweddell et al., 2002). While HLHS patients have complicated clinical course, the early mortality is largely associated with ventricular dysfunction with rapid progression to acute heart failure (HF) (Garcia *et al*., 2020). Unfortunately, therapies developed for HF in adults have been ineffective for treating HF in HLHS(Hsu et al., 2010; Shaddy et al., 2007). Without insights into the underlying mechanisms driving early HF in HLHS, the clinical management of this patient population is largely empirical.

Investigations into the mechanism of HLHS-HF have been hampered by the difficulty in obtaining human heart tissue for analysis. An alternative strategy entails in vitro disease modeling using induced pluripotent stem cells (iPSC) and their differentiated derivatives such as cardiomyocytes (iPSC-CM), endothelial/endocardial cells, and other cell types. While this has been successfully deployed for investigating HLHS disease mechanisms (Gaber et al., 2013; Hrstka et al., 2017; Jiang et al., 2014a; Kobayashi et al., 2014; Miao et al., 2020; Paige et al., 2020), no studies have explored the possibility of using iPSC-CM to model disease outcome. Particularly compelling is the question as to why only some HLHS patients develop early-HF even with the same surgical palliation, and what might be the underlying cause for HF. The feasibility to model HLHS HF in iPSC-CM is suggested by our previous studies of a mouse model of HLHS (Liu et al., 2017). We found cell autonomous defects were associated with prenatal/neonatal lethality from HF in the HLHS mutant mice. In the present study, we showed mouse iPSC-CM generated from the HLHS mice replicated defects observed in the HLHS mouse heart tissue, confirming the defects are cell autonomous and thus suitable for in vitro iPSC-CM modeling. Generating iPSC and iPSC-CM from HLHS patients dichotomized by clinical outcome, either with or without acute early-HF (Xinxiu Xu, 2018), we further investigated and demonstrated the feasibility of using patient iPSC-CM to investigate HF outcome in vitro. These studies provided surprising insights not only into possible causes for early HF in HLHS, but they also uncovered mechanisms that may protect against early HF in HLHS patients surviving heart transplant free.

## RESULTS

### Cell Autonomous Mitochondrial defects in the Ohia HLHS Mouse Model

The Ohia HLHS mouse model exhibits mid to late gestation lethality with acute heart failure characterized by severe pericardial effusion with poor cardiac contractility, this is associated with decreased proliferation and increased apoptosis (Liu *et al*., 2017). Ultrastructural analysis showed the myocardium with poorly organized thin myofilaments and altered mitochondrial morphology (Liu *et al*., 2017). Dynamic changes in mitochondria morphology play an important role in the developmentally regulated metabolic switch from glycolysis to oxidative phosphorylation, a process that also plays a critical role in regulating cardiomyocyte differentiation (Hom et al., 2011). This entails closure of the mitochondrial permeability transition pore (mPTP) and formation of a mitochondrial transmembrane potential (ΔΨ_m_) mediating oxidative phosphorylation. Using primary cardiomyocyte explants from the E14.5 *Ohia* HLHS mouse heart, we measured the ΔΨ_m,_ in cardiomyocytes from the right and left ventricle (RV, LV). A reduction was observed in both the RV and LV cardiomyocytes, indicating failure of the mPTP to close (**Figure 1A**). However, mitochondrial mass was unchanged (**Extended Data Figure S1A).**

**Figure 1.**
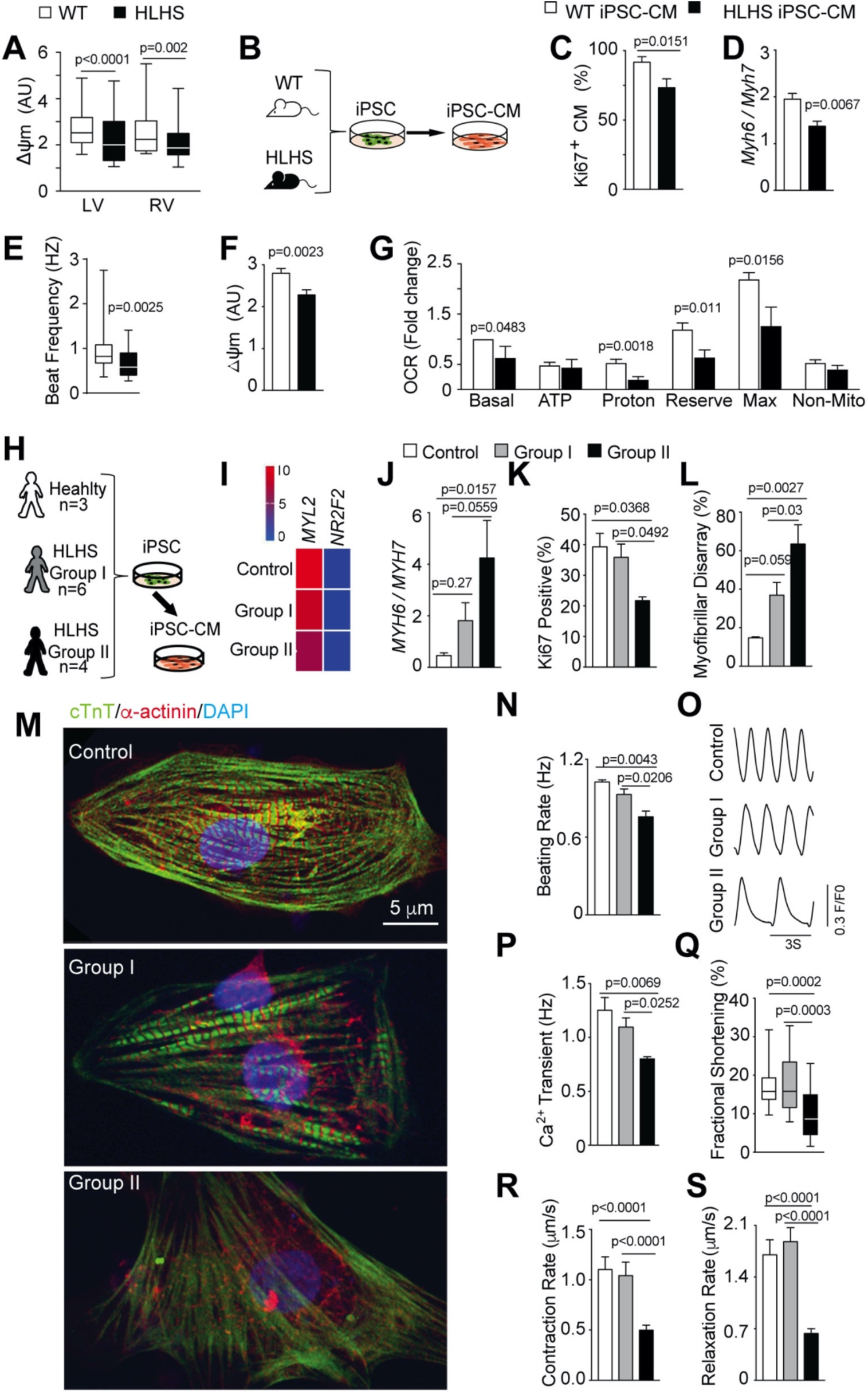
Mouse and human HLHS iPSC and iPSC-CM show differentiation and functional defects. (A) Mitochondrial transmembrane potential (ΔΨm) was measured with TMRE/Mitotracker Green in CM from E13.5 wildtype (WT) mouse embryo (n=3) left ventricle (WT-LV; n=124 CMs) and right ventricle (RV; n=86 CMs), and E13.5 *Ohia* HLHS mutant (n=3), LV (n=105 CMs) and RV (n=118 CMs) (B) Mouse iPSC were generated from WT and HLHS mouse embryonic fibroblasts (two independent lines eacg) and further differentiated into iPSC-CM. (C) Ki67 quantification showed reduced proliferation of the HLHS (n=1800) vs. WT (n=2800) iPSC-CM. (D) *Myh6*/*Myh7* transcript ratio is decreased in the *Ohia* HLHS iPSC-CM vs. WT, indicating a maturation defect. WT n=6, and HLHS n=5. (E) Beat frequency of *Ohia* HLHS iPSC-CM (n=17 clusters quantified) was reduced compared to WT (n=38 myocyte clusters quantified). (F) The mitochondrial membrane potential (ΔΨm) was reduced in the mouse HLHS iPSC-CM (n=43) compared to WT (n=56). (G) Mitochondrial respiration parameters were obtained from Seahorse Analyzer oxygen consumption rate (OCR) measurements (n=3 independent experiments). (H) Human iPSC-CM were generated from HLHS patients and controls. Group I comprises patients with transplant free survival >5 years old. Group II are patients who died or survived with heart within one year of age. Controls are healthy subjects without disease. Functional assessments were conducted on 18-22 days of iPSC-CM differentiation. (I) qPCR for atrial (NR2F2) and ventricle MYL7) marker genes show the iPSC-CM are ventricle-type. (J) *MYH6/MYH7* transcript ratio is increased in Group II iPSC-CM, indicating cardiomyocyte maturation defect. Note this ratio is reversed in mice vs. human, as the major ventricular myosin heavy chain in mice is *Myh6*, and *MYH7* in human. (K) Ki67 immunostaining showed decreased proliferation in Group II iPSC-CM. (L, M) Quantification of myofibril organization showed myofibrillar disarray (L) in cTnT (Green) and α-actinin (Red) positive human iPSC-CM (M). (N-P) Beat frequency (N), and visualization (O) and quantification of calcium transients (P) in iPSC-CM showed functional deficits in Group II iPSC-CM. (Q-S) Quantification of contractile function in individual iPSC-CM showed decreased fractional shortening (Q), contraction rate (R) and relaxation rate (S) in Group II iPSC-CM. Data shown are mean±SEM using Student’s t-test or ANOVA. For box plots, median/min/max are shown with Kruskal-Wallis statistics. Number of Control, Group I, Group II subjects analyzed (H-S): (I) n=3,3,3 subjects. (J) n=3,5,3 subjects. (K, L,N) n=3,6, 3 subjects. (P) n=3,6,4 subjects. (Q-S) n=3,4,4 with n=17,23,38 cardiomyocytes respectively.

To determine whether the abnormal open state of the mPTP is a cell autonomous defect, we generated iPSC from *Ohi*a mutant fibroblasts and differentiated them into iPSC-CM (**Figure 1B**). These *Ohia* iPSC-CM generated entirely *in vitro* showed reduced cell proliferation with lower *Myh6/Myh7* ratio indicating a cardiomyocyte differentiation defect (**Figure 1C,D**;Figure S1C,D), phenotypes reminiscent of those observed in the *Ohia* HLHS myocardium. Poor cardiac function was also indicated by reduced beat frequency (**Figure 1E**). Mitochondrial function was assessed with measurement of ΔΨ_m_ and oxygen consumption rate (OCR) using the Seahorse Flux Analyzer (**Figure 1F,G**). This analysis uncovered mPTP and mitochondrial respiration defects in both the undifferentiated *Ohia* iPSC and iPSC-CM. While the iPSC showed lower respiratory reserve and respiratory maxima (Figure S1B), the iPSC-CM from *Ohia* exhibited reduction in basal respiration, ATP production, respiratory reserve, and respiratory maxima (**Figure 1G**; Figure S1E). Together these findings indicate the mitochondrial dysfunction, and proliferation and differentiation defects observed in the Ohia HLHS heart tissue are cell autonomous.

### Generating HLHS Patient iPSC-CM for Investigating Early Heart Failure

The finding that *Ohia* iPSC-CM replicated defects seen in the HLHS heart tissue suggested HLHS patient derived iPSC-CM may have utility for investigating acute early HF in HLHS patients. For this study, we generated iPSC from 10 HLHS patients, including six >5-year old with transplant free survival (Group I) (**Figure 1H;Figure S2A**), and four that died (n=3) or survived (n=1) with a heart transplant at <1 year of age (Group II). In addition, we also generated iPSC from 3 healthy subjects as controls. Using standard iPSC-CM differentiation protocols, iPSC-CM at Day 16-20 of differentiation were generated and used for the subsequent analysis.

### Impaired Cardiomyocyte Differentiation and Contractile Dysfunction

The iPSC-CM were found to be predominantly ventricle-like as shown by high expression of the ventricular marker MYL2, but low expression of atrial marker NR2F2 (Biendarra-Tiegs et al., 2019) (**Figure 1I**). The Group II iPSC-CM had fewer cardiac troponin T (cTnT) positive cells with higher ratio of *MYH6* (atrial myosin heavy chain) to *MYH7* (ventricular myosin heavy chain) transcripts, indicating poor differentiation (Jiang et al., 2014b) (**Figure S2E;Figure 1J**). Group II iPSC-CM also showed reduced Ki67, but increased pH3 immunostaining, suggesting cell cycle disturbance with possible metaphase arrest (**Figure 1K**; Figure **S2F,G**), reminiscent of findings in the *Ohia* HLHS LV (Liu *et al*., 2017). Poor cardiomyocyte differentiation was indicated by low expression of cTnT (A-band) and α-actinin (Z-disc) containing myofilaments together with increased myofibrillar disarray (**Figure 1L, M**). However, no change was observed for cardiomyocyte cell size (**Figure S2H**).

Further assessment of cardiomyocyte contractile function showed the Group II iPSC-CM have lower beat frequency with reduced calcium transients (**Figure 1N-P;Supplemental Video 1&2**). Examination of the profile of calcium transients confirmed the majority (84∼89%) of the iPSC-CM are ventricle-like (Cyganek et al., 2018) (**Figure S2I)**. Analysis of the cardiomyocyte contractile motion by high resolution video microscopy showed reduced fractional shortening accompanied by decreased contraction and relaxation rates in the Group II but not Group I iPSC-CM. This was associated with reduction in the diastolic sarcomere length, but not systolic sarcomere length **(Figure 1Q-S;Figure S2J,K; Supplemental video 3)**. Together these findings indicate the Group II iPSC-CM have profound differentiation defects causing impaired calcium handling and poor contractile function.

### Mitochondrial Respiration and Transition Pore Closure Defects

Various parameters of mitochondrial function were assessed in the iPSC-CM. A marked decrease in mitochondrial membrane potential (ΔΨ_m_) was observed in the Group II iPSC-CM, suggesting abnormal mPTP opening (**Figure 2A**). OCR measurements showed mitochondrial respiration defects with reduction in basal respiration, ATP production, H+ leak, respiratory reserve and maximal respiratory capacity (**Figure 2B**). For Group I iPSC-CM, only respiratory reserve and maximal respiratory capacity showed significant change (**Figure 2B**). These same two parameters also were reduced in the undifferentiated iPSC of Group II patients, findings reminiscent of the *Ohia* mouse iPSC (**Figure S2C & S1B**). Western blotting showed no change in abundance of the electron transport chain (ETC) complexes **(Figure S3A,B)**.

**Figure 2.**
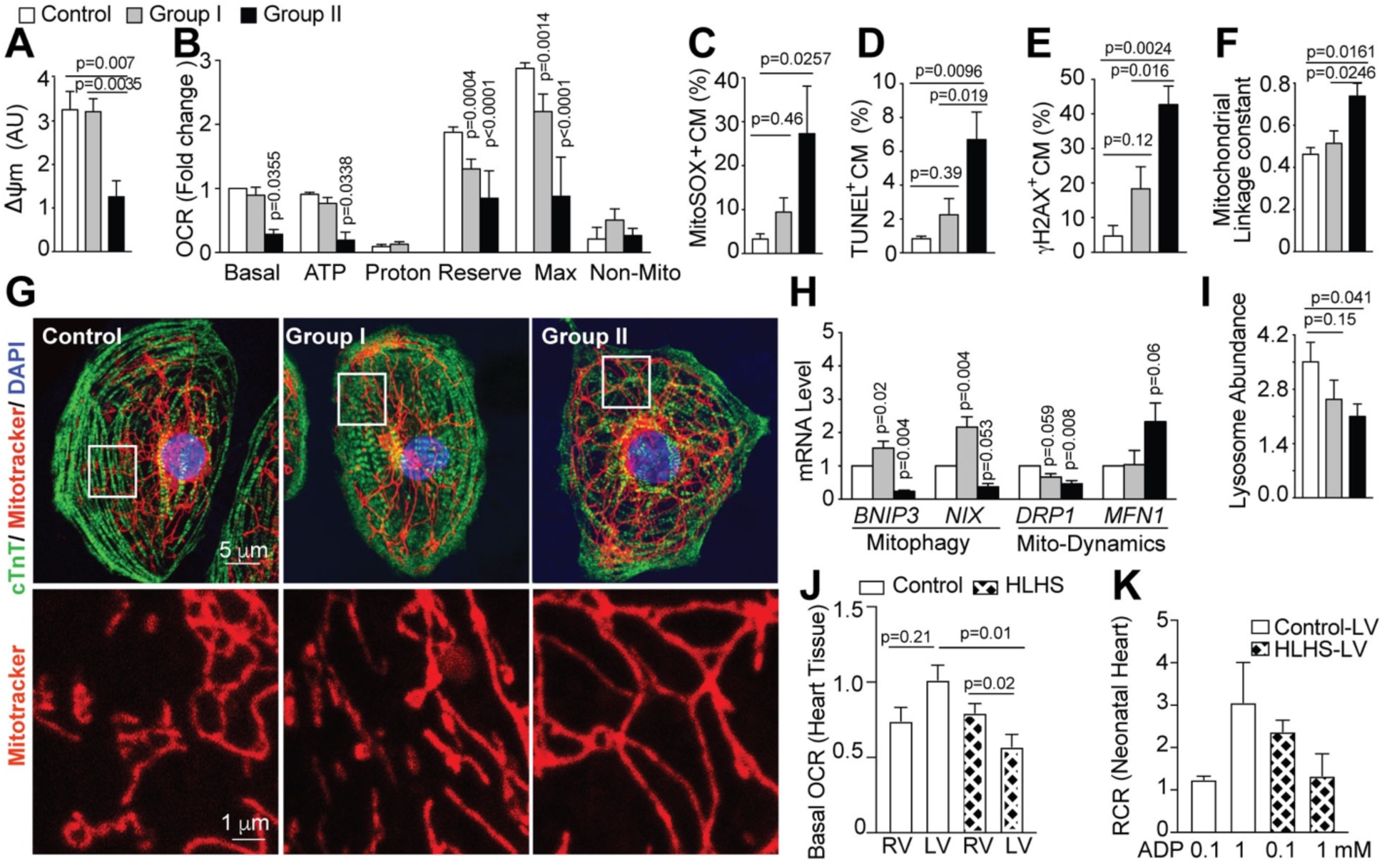
Mitochondrial dynamics and respiration defects in HLHS iPSC-CM. (A) Measurement of ΔΨm in human iPSC-CM using TMRE and MTG staining. (B) Seahorse Analyzer OCR measurement showed mitochondrial respiration defects in the HLHS iPSC-CM. (C) MitoSOX staining show elevated mitochondrial reactive oxygen species (ROS) in Group II HLHS iPSC-CM (D,E) TUNEL labeling and *γ*-H2AX staining show increased apoptosis (D) and DNA damage (E) in Group II iPSC-CM. (F,G) Mitotracker red staining showed increased linkage constant (see Methods), indicating hyperfused mitochondria in Group II iPSC-CM (H) qPCR of key genes regulating mitophagy and mitochondrial dynamics. (I) LysoTracker Deep Red-staining of lysosomes showed lysosome reduction in Group II iPSC-CM. (J) Basal OCR (>3 duplicates) of explanted heart tissue from two 19-year-old HLHS patients and a 15 year old cardiomyopathy patient with doxorubicin cardiotoxity were assessed using the Seahorse Analyzer (K). Respiratory control ratio (RCR) was obtained for 2 Control and 2 HLHS neonatal patient using cell extracts from explanted heart tissue. V_max_ was measured using succinate as a substrate and two concentrations of ADP. Bar graphs show mean±SEM with ANOVA. Number of Control, Group I, Group II subjects analyzed respectively in (A-I): (A, D) n=3,6,3 subjects. (B, C,F) n=3,5,3 subjects,(E) n=3,4,3 subjects. (H) n=3,3,3 subjects (I) n=3, 4, 3 subjects.

Consistent with the uncoupling of oxidative phosphorylation in the Group II iPSC-CM, we observed a marked increase in mitochondrial reactive oxygen species (ROS) indicated by increased MitoSOX staining (**Figure 2C**) (Hom *et al*., 2011). Also observed was a reduction in nitric oxide (NO), suggesting perturbation of protein nitrosylation required for normal mitochondrial respiration (**Figure S3F**). The mitochondrial respiration defects and increase in ROS in the Group II iPSC-CM are accompanied by increase in apoptosis and activation of a DNA damage response, findings similar to those observed in *Ohia* (Liu *et al*., 2017) and human HLHS fetal heart tissue (Gaber *et al*., 2013) (**Figure 2D-F**). Parallel analysis of the Group I iPSC-CM showed no significant change in these parameters.

### Perturbation of Mitochondrial Dynamics

As the regulation of mitochondrial fission and fusion play important roles in metabolic and redox regulation, its disturbance can contribute to cardiomyocyte death in HF (Marin-Garcia and Akhmedov, 2016). Using confocal imaging, we assessed mitochondrial mass and morphology. In the Group II but not Group I iPSC-CM, mitochondrial linkage constant and cluster length were increased, while mitochondrial mass was unchanged, indicating increase in mitochondrial fusion in the Group II iPSC-CM (**Figure 2F,G; Figure S3C-E).** Group II iPSC-CM also showed decreased expression of *DNML (DRP1),* gene regulating mitochondrial fission, and increase in *MFN1,* gene promoting mitochondrial fusion (**Figure 2H**). Expression of *BNIP3*/*NIX* regulating mitophagy were decreased in Group II but increased in Group I iPSC-CM (**Figure 2H**). Lysosomes, which are involved in mitophagy, were reduced in both Group I and II (**Figure 2I**). These findings indicate the hyperfused mitochondria in the Group II iPSC-CM likely arise from altered mitochondrial dynamics associated with increased mitochondrial fusion and decreased mitophagy. In contrast, Group I iPSC-CM exhibited more normal mitochondrial dynamics that may be accompanied by increase in mitophagy.

### Mitochondrial Respiration Defects in the Left Ventricle of HLHS Human Heart Tissue

To explore the clinical relevance of the abnormal mitochondrial function observed in the HLHS patient derived iPSC-CM, heart tissue was obtained from HLHS patients undergoing heart transplant. Analysis of mitochondrial respiration showed basal respiration was reduced in the HLHS-LV vs. RV tissue, but such LV-RV difference was not observed in heart tissue from age-matched heart transplant patient with doxorubicin induced HF (**Figure 2J**). Analysis of two additional HLHS neonates and two neonatal control subjects the HLHS-LV being more sensitive to lower ADP, indicating possible adaptation to bioenergetic stress. However, respiration in the LV failed to increase with increasing ADP concentration, indicating the hypoplastic LV has reduced respiratory capacity (Ventura-Clapier et al., 2011) (**Figure 2K;Figure S3G**). Western blotting showed no change in ETC components (**Figure S3H,I**). These findings suggest LV specific mitochondrial respiration defects in HLHS.

### Defects in Yap-Regulated Antioxidant Response

Activation of an antioxidant defense pathway occurs during developmental with metabolic transition to mitochondrial respiration (Perrelli et al., 2011; Tsutsui et al., 2011). This pathway is regulated by transcription factors NRF2 (Itoh et al., 1999) together with PITX2 and YAP (Tao et al., 2016). These three transcription factors play an essential role in regulating the expression of antioxidant genes that scavenges ROS to prevent oxidative stress. These transcription factors also have critical roles in regulating cardiac regeneration and repair, with YAP also shown to regulate heart organ size (Heallen et al., 2013; von Gise et al., 2012; Zhou et al., 2015). Interestingly, YAP also has been demonstrated to regulate mitochondrial fission (Huang et al., 2018).

We observed *NRF2* and *PITX2* transcripts are both reduced in Group II iPSC-CM, but in Group I iPSC-CM, *PITX2* was elevated and *NRF2* was unchanged (**Figure 3A**). In contrast, *YAP1* transcripts showed no change in either Group or Group II iPSC-CM (data not shown). Analysis of downstream genes in the antioxidant pathway revealed up regulation of thioredoxin (*TXN*), peroxiredoxin 1 (*PRDX1*), glutathione peroxidase 1 (*GPX1*), and superoxide dismutase 2 (SOD2) in the Group I iPSC-CM, but in Group II, expression was either unchanged or downregulated, such as for PRDX1 (**Figure 3A**). In the Group I iPSC-CM, we also observed increased expression of HIF1α, a transcription factor regulating cell stress response to hypoxia. This was associated with increased expression of *VEGF*, a known downstream transcriptional target of HIF1α (**Figure 3A**) (Guimaraes-Camboa et al., 2015).

**Figure 3.**
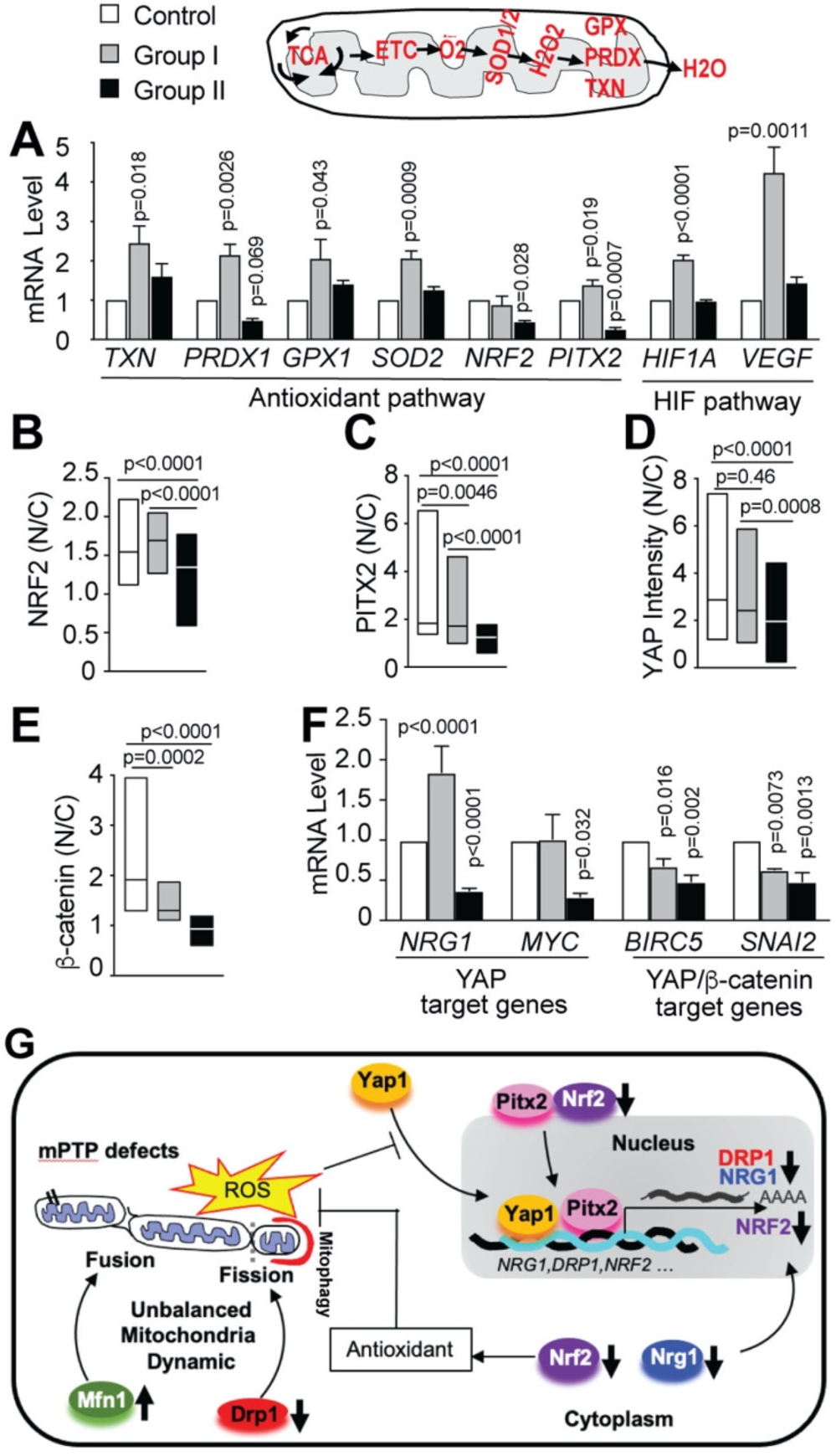
HLHS iPSC-CM with failed antioxidant response show cytoplasmic localization of NRF2, YAP1, and PITX2. (A) qPCR showed key antioxidant genes and HIF pathway are up regulated in Group I iPSC-CM, and either unchanged or downregulated in Group II iPSC-CM. (B-E) Immunostaining show defect in NRF2, PITX2, YAP and β-catenin nuclear localization in Group II iPSC-CM (see FigS6 G). N/C = nuclear to cytoplasmic ratio. (F) qPCR quantification of YAP and YAP/β-actinin downstream target genes. (G) Diagram summarizing HLHS associated defects in mitochondrial dynamics with elevated ROS and failed antioxidant response with failure in NRF2, YAP, and YAP/PITX2 nuclear translocation. (A, F) show mean±SEM with one-way ANOVA. Box plot with median/min/max shown with Kruskal-Wallis statistics. n=3 control, 3 Group I, 3 Group II subjects. Number of Control, Group I, and Group II subjects analyzed: (B, C) n=130, 117, 64 CM. (D) n=90, 78, 79 CM. (E) n=44, 15, 17 CM

Antibody staining showed nuclear localization of NRF2/PITX2/YAP were reduced in the Group II iPSC-CM, while in Group I, only PITX2 showed a modest reduction in comparison to Group II and control (**Figure 3B-D**; **Figure S3J**). However, total YAP and β-catenin protein expression levels were unchanged (**Figure S3K,L**). We further examined expression of downstream target genes of YAP - NRG1 and MYC *(Artap et al., 2018)*, and observed both were reduced in the Group II iPSC-CM. In contrast, the opposite was observed in Group I with *NRG1* being upregulated, while *MYC* was unchanged (**Figure 3F**). However, nuclear localized β-catenin and transcripts for two downstream YAP/β-catenin targets, *BIRC5* and *SNAI2*, were reduced in both Groups I and II iPSC-CM (**Figure 3E,F**). Together these findings show defects in the mounting of an effective antioxidant response in the Group II iPSC-CM (**Figure 3G**). In contrast, in Group I, the antioxidant capacity is expanded, and may promote and support the restoration of redox homeostasis.

### Inhibition of mPTP Opening Rescues Mitochondrial Respiration and Apoptosis

The mPTP closure defect observed in the Group II iPSC-CM suggests compounds promoting mPTP closure might rescue the mitochondrial defect. This was investigated using the Seahorse Analyzer to screen compounds known to inhibit mPTP opening(Martel et al., 2012) or otherwise modulate mitochondrial respiration. This analysis yielded 7 compounds showing some rescue of mitochondrial respiration in the Group II iPSC-CM. This included fasudil, sildenafil, cyclosporin A (CsA), Metformin, JP4, SS31, and Y27632 (**Figure S4A**). In contrast, treatment with ascorbic acid, a general antioxidant, had no effect (Myung et al., 2013; Ye et al., 2013). Given sildenafil is commonly used among CHD patients for its vasodilatory effects(Galie et al., 2005), more in-depth analysis was carried out with sildenafil (**Figure 4**).

**Figure 4.**
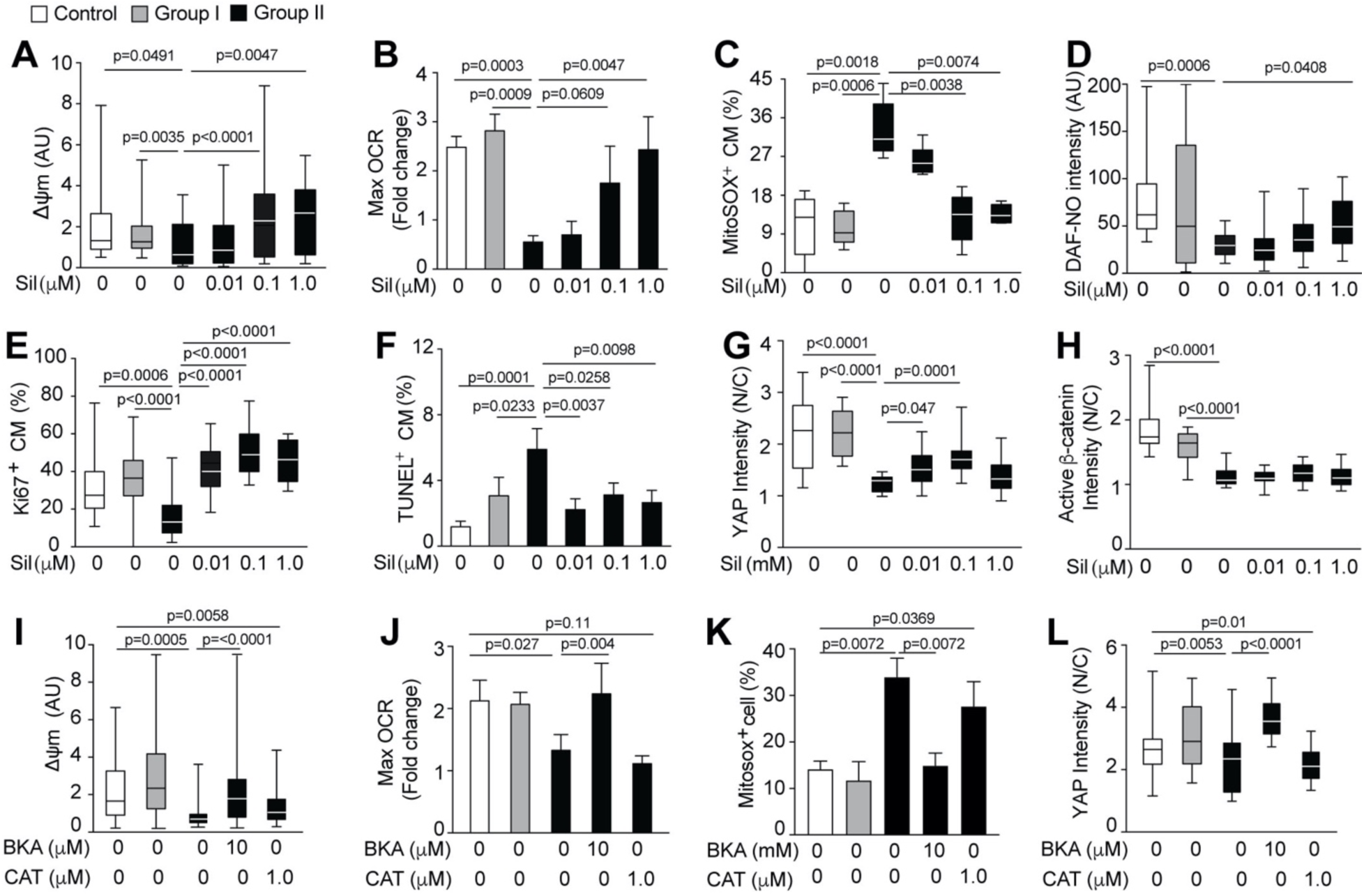
Inhibition of mitochondrial membrane permeability rescues mitochondrial respiration and apoptosis. (A-H) Sildenafil (Sil) rescued Group II iPSC-CM including ΔΨm (A), maximum OCR (B), mitochondrial ROS (C), NO level (D), cell proliferation (E), apoptosis (F), and YAP (G) and β-catenine (H) nuclear localization. (I-L) Treatment with bongkrekic acid (BKA), but not carboxyatractyloside (CAT) rescued ΔΨm (I), maximum OCR (J), mitochondrial ROS (K) and YAP nuclear localization (L). Bar graphs show mean±SEM, analyzed by one-way ANOVA, and box plot with median/min/max analyzed by Kruskal-Wallis. n≥3 independent repeats. Subjects analyzed: Control n=3, Group I n=4 or 5, and Group II, n=3 or 4.

Titration of sildenafil showed rescue down to 0.1 μM, which restored not only ΔΨ_m_ and maximal mitochondrial respiration. This also reduced mitochondrial ROS to levels similar to the Group I and control iPSC-CM (**Figure 4A-C; Figure S4B-D**). Sildenafil is also known to affect NO production, but normal NO level was restored only at 1.0 μM concentration (**Figure S4D**)(Prabhu et al., 2013). Cell proliferation, apoptosis (**Figure 4E,F**), and YAP nuclear trafficking were rescued at ten times lower dose of 0.01 μM (**Figure 4G**). However, β-catenin nuclear trafficking was not rescued (**Figure 4H; Figure S4E**). To verify that sildenafil is targeting the mPTP, we further assessed treatment with BKA (bongkrekic acid) and CAT (carboxyatractyloside), which activate and inhibit the mPTP, respectively. As expected, BKA but not CAT rescued the mPTP defect, with opposing effects observed for maximal and basal OCR, mitochondrial ROS, and YAP nuclear localization (**Figure 4I-L, Figure S4F**). Similar treatment of control iPSC-CM showed repression of respiration by CAT, while BKA had no effect (**Figure S4G**).

### Single Cell Transcriptome Profiling

We performed single cell RNAseq on iPSC-CM from two Group II patients, 7042 with heart transplant at 11 months and patient 7052 deceased at 2 months, Group I patient 7464 surviving transplant free at 7 years of age, and healthy control subject 1053. Data was obtained from 4403 cardiomyocytes forming 9 clusters (Stuart et al., 2019) (**Figures 5A and S5A-C**). Marker gene analysis showed these cardiomyocytes were largely of ventricular identity (**Figure S5D)**. Clusters 0 (CM I), 1 (CM II) and 5 (CM III) comprising the majority of cells are well differentiated cardiomyocytes of increasing maturation (**Figure S5E; Supplemental Spreadsheet 2**). Group II vs. control comparison yielded the greatest number of DEGs (**Figure 5B**). Enrichment was observed for mitochondrial related pathways in all three clusters, suggesting Group II mitochondrial defects likely arise early in cardiomyocyte differentiation (**Figure 5C**). In contrast, Group I vs Control yielded the fewest DEGS. These were associated with heart development and muscle organ development terms in Clusters 0 and 1, and mitochondrial related terms in Cluster 5 (**Figure 5E**). Group II vs. Group I comparisons yielded apoptosis and cell death in Clusters 0 and 1, (**Figure 5D**), and tRNA modification and noncoding RNA in Cluster 5.

**Figure 5.**
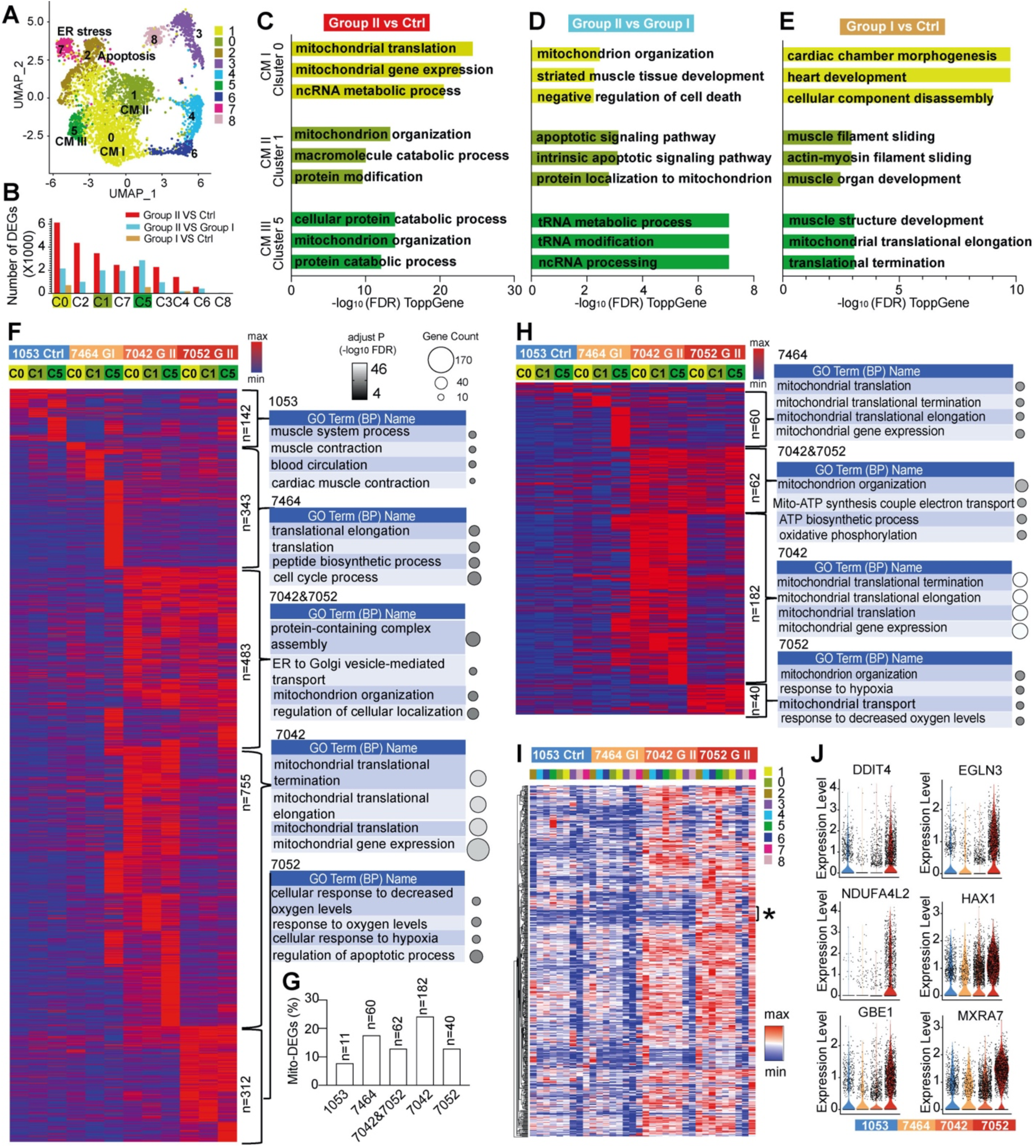
Single cell RNAseq showed mitochondrial pathways associated with early-HF. (A) Single cell RNAseq data yielded 9 distinct clusters (See Figure S5E). Clusters 0,1, and 5 are comprised of well differentiated cardiomyocytes of increasing maturation. Clusters 4 and 6 correspond to proliferating cardiomyocytes at G2/M and S phase (also see Figure S5E). Cluster 2 include cardiomyocytes undergoing apoptosis, while Cluster 7 exhibit evidence of ER stress with UPR (Figure S5E). In Clusters 3 and 8, enrichment for oxidative phosphorylation and genes related to hypertrophic cardiomyopathy are observed (see Spreadsheet 2). (B) DEGs recovered in each cluster with pairwise comparisons. (C-E) Pathway enrichment of with DEGs in pairwise patient group comparisons in Clusters 0, 1 and 5. (F). Heat map of DEGs from all pairwise comparisons for Clusters 0,1, and 5 and the top Biological Processes recovered. (G). Percentage of DEGs that are mitochondrial related is shown for each group in F. The n represents number of mitochondrial DEGs observed. (H). Heat map of mitochondrial related DEGs in clusters 0, 1 and 5 and the top Biological Processes recovered. (F, H) Color scale showed relative maximum and minimum value. Grayscale showed the adjust p value (-log10FDR) of each GO term and circle size showed the count of genes in each GO term. (I) Hierarchical clustering based on DEGs from Group II (7052 and 7042) vs. Group I (7464). Asterisk (*) denotes region with 28 DEGs upregulated only in 7052 (Spreadsheet 2). (J) Violin plot showing transcript expression for 6 of the 28 DEGs from panel I.

Combining DEGs from all pairwise comparisons showed the number of DEGs increased with disease severity (**Figure 5F**). Only control 1053 yielded terms related to muscle and muscle contraction. Group I 7464 recovered protein translation and cell cycle, and Group II 7042 yielded mitochondria and mitochondrial translation. In Group II 7052, hypoxia related pathways were recovered, but mitochondrial related terms were also recovered in DEGs shared with 7042 (**Figure 5F; Spreadsheet 2**). Overall, a high percentage of the DEGs were found to be mitochondrial related (**Figure 5G**). Heatmap generated comprising only the mitochondrial related DEGS was nearly identical to that for all DEGs (**Figure 5H** **vs. F**), indicating genes with the highest fold change are mostly mitochondrial related. Interestingly pathways related to mitochondrial translation, elongation, and termination were recovered in both 7464 (Group I) and 7042 (Group II), but with only 60 DEGs in 7464, vs. 183 DEGs in 7042 (**Figure 5H;Spreadhseet 2**). Differing from 7042, mitochondrial DEGs in 7052 were hypoxia related, confirming the recovery of these same pathways in all DEG analysis (**Figure 5F**). In mitochondrial DEGs shared between 7042/7052 shared, the recovery of ATP synthesis and oxidative phosphorylation were observed, suggesting bioenergetic deficits associated with Group II patients.

DEG analysis based on Group II vs. Group I comparison yielded further evidence of the effective dichotomization of HLHS patients into two functional groups (**Figure 5I**). Profiling upregulated DEGs showed the two Group II patients are similar to each other, while Group I is similar to control (**Figure 5I**). This analysis also recovered 28 genes highly expressed only in patient 7052 (see region denoted by asterisk in **Figure 5I**) - 14 are related to mitochondria, hypoxia and/or cell death, including EGLN3 encoding prolyl hydroxylase, an oxygen sensor that promotes HIF1α degradation (**Figure 5J;Supplemental Spreadsheet 2**). Assembly of a protein interactome network incorporating 26 of these genes showed pathway enrichment for hypoxia, apoptosis, and oxidative stress, indicating these genes are part of a functional network contributing to early HF in patient 7052 (**Figure S6**;**Supplemental Spreadsheet 2**).

### Molecular Chaperone Rescues Mitochondrial Respiration and Apoptosis

Recovery of ER stress and UPR in Cluster 7 from the scRNAseq analysis was notable (**Figure 6A**), given ER stress can be triggered by mitochondrial dysfunction and oxidative stress. and ER stress has been associated with HF (Schiattarella et al., 2019). This pathway has not been investigated previously in the context of HLHS. Real time PCR analysis confirmed elevated expression of genes (*XBP1,ATF4,ATF6*) associated with the three conserved ER stress pathways (**Figure 6B**). All three pathways were elevated in patient 7052, and two (*ATF4 ATF6*) were elevated in 7042. In contrast, Group I patient 7464 showed no change relative to control (**Figure 6B**). Similar analysis of three downstream ER stress target genes *HSPA5, DDIT3,* and *DNAJC3* showed all three were upregulated in 7052, but only *DDIT3* and *DNJC3* were elevated in 7042. In contrast, all three genes were down regulated in Group I patient 7464 (**Figure 6B**). We noted these same cell stress related genes were also up regulated in the *Ohia* HLHS heart tissue, consistent with their severe HF phenotype (Liu *et al*., 2017). To assess the potential functional impact of UPR on the HLHS iPSC-CM, we treated the iPSC-CM with Tauroursodeoxycholic acid (TUDCA), a molecular chaperone known to promote protein folding and suppress ER-stress. TUDCA treatment promoted mPTP closure, reduced mitochondrial ROS, and rescued NO production in the Group II iPSC-CM (**Figure 6C-E**). TUDCA also rescued YAP nuclear translocation, restored cardiomyocyte proliferation and blocked apoptosis (**Figure 6F-H**). These findings suggest ER stress and UPR may contribute to the early HF in Group II patients.

**Figure 6.**
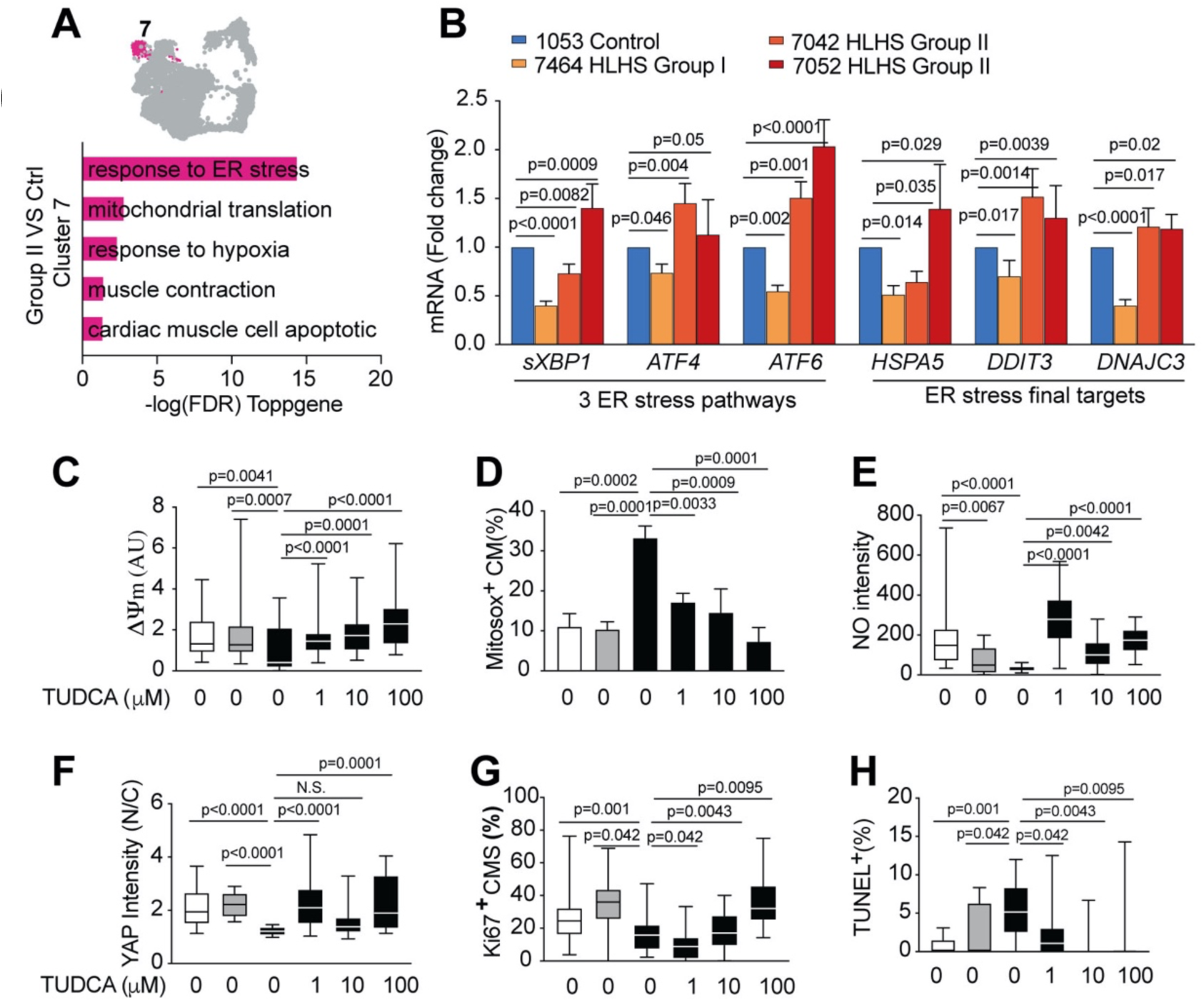
ER stress in the iPSC-CM and its suppression rescued mPTP closure and apoptosis. (A) ER stress was recovered as top pathway in Cluster 7. (B) Real time PCR confirmed elevated expression of ER stress marker genes in Group II (7042,7052) iPSC-CM. (C-H) Treatment with molecular chaperone TUDCA, an ER stress inhibitor, rescued ΔΨm (C), ROS (D), NO level (E). YAP nuclear localization (F), and restored cell proliferation (G) and suppressed apoptosis (H) Bar graphs show mean±SEM with Student’s t-test. Box plots show median and minimum-maximum, with Mann-Whitney statistical test. (B) n=3 independent repeats for each sample. (C-H) n≥3 independent repeats for each bar. Subjects analyzed: Control n=3, Group I n=4 or 5, and Group II, n=2 (7042 and 7052).

### Enrichment of variants associated with mitochondrial metabolism

Given HLHS is well described as having a genetic etiology, we further investigated the whole exome sequencing data available for 6 of our 10 HLHS patients (Group II 7042, Group I 7131,7400,7438,7434,7464). High impact variants comprising unique loss-of-function variants were recovered (**Figure S7A**;S**upplemental Spreadsheet 3**). Using Webgestalt/KEGG pathway enrichment analysis, four genes were identified as significantly associated with metabolic pathways (*OXA1L,NNMT,NEU3,ALDH7A1*) (S**upplemental Spreadsheet 3**). An protein-protein interactome (PPI) was constructed using these four genes to explore connections to Hippo signaling, a pathway that plays a critical role in regulating YAP degradation and nuclear translocation (Meng et al., 2016). The interactome showed enrichment for Hippo and Wnt signaling, and also heart development and many mitochondrial-related terms, including regulation of mitochondrial membrane permeability (**Fig7A,B;**S**upplemental Spreadsheet 3**).

Building on this finding, we interrogated the WES data from another 41 HLHS patients comprising 19 patients who died or had heart transplant (unfavorable outcome) and 22 HLHS patients surviving transplant-free beyond 5 years of age (favorable outcome). Interrogating for unique loss-of-function variants or predicted damaging missense or splice variants yielded 159 genes from the unfavorable outcome group and 194 genes from the favorable group (**Figure S7A;**S**upplemental Spreadsheet 3**). Rendering these genes in a network plot using Metascape recovered terms such as “Ion channel transport”, “Mitochondrial gene expression”, and “Mitochondrial translation” in association with genes from the unfavorable group, while “lipid location, response to IL-17” were associated with the favorable group (**Figure 7C**;S**upplemental Spreadsheet 3)**. Some pathways were shared by both groups such as calcium signaling, MAPK signaling, and nervous system development. Examining the genes recovered for intersection with an expanded MitoCarta-related inventory of mitochondrial genes yielded 19 genes from the unfavorable and 11 from favorable group (Calvo et al., 2012; Pagliarini et al., 2008) (S**upplemental Spreadsheet 3**). ToppGene analysis of these overlapping genes recovered from the unfavorable group yielded multiple mitochondrial related pathways, including mitochondrial translation (**Figure 7D**). Most of these genes are highly expressed in cardiomyocytes of the human fetal heart (Cui et al., 2019), supporting a role in HLHS pathogenesis (**Figure 7E**).

**Figure 7.**
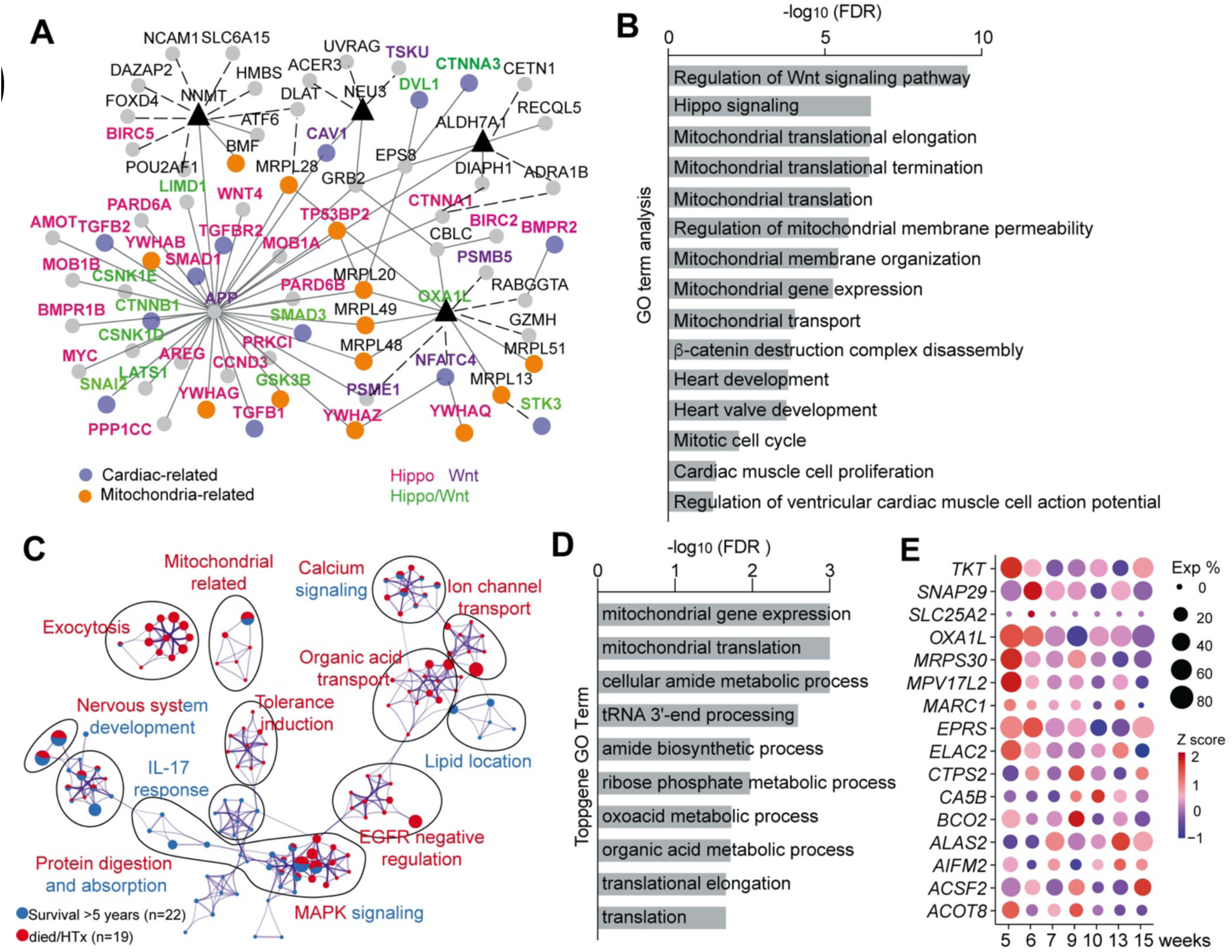
Damaging variants in mitochondrial and Hippo related pathways. (A,B) Protein-protein interactome constructed with four genes (*OXA1L, NNMT, NEU3, ALDH7A;* blue triangle nodes) to explore interconnections with the Hippo pathway recovered many genes related to Hippo and Wnt signaling and also mitochondrial and cardiac related genes (C) Pathway enrichment rendered using Metascape comprising genes recovered with extreme unique variants from 41 HLHS patients. Size of the collective node slice represents percentage of genes originating from the corresponding gene list. (D) Pathway enrichment related to the 19 mitochondrial genes recovered from the unfavorable outcome HLHS patients that intersected with the MitoCarta inventory of mitochondrial genes (see overlapping genes in Supplementary Spreadsheet 3). (E) Mitochondrial transcript expression in cardiomyocytes from human heart tissue at 5-15 weeks gestation from scRNAseq data of Cui, et al (Cui *et al*., 2019). Size of the circle corresponds to percentage of cells expressing the gene (Exp%) and the color show average expression values with Z-transform.

## DISCUSSION

Our objective in this study was to investigate why only some HLHS patients suffer early-HF and the possible cause of early-HF. Analysis of iPSC-CM from our HLHS mouse model and HLHS patients revealed both have cell autonomous defects involving failure in mPTP closure. Thus, mitochondrial defects seen in vivo in the HLHS mouse heart were replicated in the mouse iPSC-CM. This was associated with defects in mitochondrial respiration and poor cardiomyocyte differentiation. The mitochondrial defects observed in the mouse heart and iPSC-CM were replicated in iPSC-CM of HLHS patients with early HF, suggesting a common cell autonomous mechanism involving mitochondrial defects underlying early-HF in HLHS.

For these studies, we selected HLHS patients with extreme phenotype comprising death or surviving with heart transplant at less than one year of age (Group II), as the first year of life poses the greatest risk with 30% mortality reported (Oster *et al*., 2013). For comparison, HLHS patients surviving transplant free at more than 5 years of age were recruited (Group I). Using these two HLHS group comparisons and control subjects, we interrogated a myriad of parameters such as cardiomyocyte differentiation, myocyte contractility and calcium handling, mPTP closure, mitochondrial dynamics, respiration, and regulation of the antioxidant pathway. From this comprehensive analysis, we showed the iPSC-CM from the Group II patients closely resembled each other, while the Group I patients were more similar to control. This was further corroborated with scRNASeq analysis, which showed transcriptome profiles of the two Group II patients are similar to each other, but very different from Group I patient 7464.

In the Group II iPSC-CM, we uncovered severe oxidative stress arising from mitochondrial dysfunction. This is associated with mPTP closure defect with altered mitochondrial dynamics and reduced mitophagy. When combined with a failed antioxidant response, this would exacerbate the redox stress to enhance apoptosis and increase DNA damage. Also observed were severe defects in cardiomyocyte differentiation with poor myocyte function. Cardiomyocyte differentiation and maturation are known to be regulated by mPTP closure (Hom *et al*., 2011) and a metabolic switch to mitochondrial respiration (Mills et al., 2017; Nakano et al., 2017) . We note skeletal myoblast differentiation is also regulated by a similar metabolic transition(Fortini et al., 2016a). Moreover, this skeletal myoblast metabolic transition was shown to be modulated by mitophagy(Fortini et al., 2016b). Also observed in the Group II iPSC-CM is the up regulation of ER stress pathways. This likely occurs secondary to the mitochondrial associated increase in ROS, exacerbating the oxidative stress induced apoptosis. Recent studies have in fact shown an important role for ER stress and UPR in HF (Schiattarella *et al*., 2019). Of significant interest from a therapeutic standpoint, apoptosis in the Group II iPSC-CM can be rescued using sildenafil (Ascah et al., 2011) to inhibit mPTP opening or TUDCA to suppress UPR. This was associated with the reduction of mitochondrial ROS, recovery of mitochondrial respiration, and restoration of YAP nuclear translocation. Together these findings support mitochondrial mediated oxidative stress as underlying the acute early-HF in HLHS. The scRNAseq analysis further suggests this may involve defects in the HIF1α pathway, altered mitochondrial translation, and bioenergetic deficits, findings that will need to be further investigated in future studies.

In contrast to Group II iPSC-CM, the Group I iPSC-CM show similarities to that of control with near normal mitochondrial respiration and normal mitochondrial dynamics without oxidative stress nor increase in apoptosis. Nevertheless, the Group I iPSC-CM have reduced mitochondrial respiratory reserve and reduced maximal respiration, indicating an overall reduction in total respiratory capacity. Importantly, nuclear localization of NRF2, YAP1, PITX2 was maintained, albeit with some reduction observed for PITX2. This was associated with striking gene expression changes that included elevated expression of many antioxidant genes, and the elevated expression of HIF1α and its downstream target genes. Genes regulating mitophagy were also elevated, while *MFN1*, gene regulating mitochondrial fusion was down regulated. Significantly, key mediators of all three ER stress pathways were downregulated. Together these findings suggest the maintenance of mitochondrial dynamics in conjunction with the suppression of oxidative and ER stress by a vigorous NRF2/YAP/PITX2 mediated antioxidant response may provide protection from early-HF in Group I patients. As Group I iPSC-CM also showed better differentiation with improved myocyte contractile function, these factors also may contribute to improved clinical outcome.

The WES sequencing analysis showed pathogenic variants in HLHS patients with unfavorable outcome are enriched for genes in mitochondrial related pathways. While the genetic causes for HLHS remains largely unknown, pathogenic variants in mitochondrial related pathways may contribute to the pathogenesis of HLHS or they may act as genetic modifiers affecting clinical outcome. It is worth noting *Sap130*, one of the two genes causing HLHS in the *Ohia* mouse model is known to regulate genes involved in mitochondrial metabolism via the Sin3A complex (Pile et al., 2003), suggesting the developmental etiology of HLHS may involve the disturbance of mitochondrial metabolism (Liu *et al*., 2017). We note there is mounting evidence of the integral role for metabolism and mitochondrial respiration in the regulation of a wide range of developmental processes (Mills *et al*., 2017).

In summary, our findings point to the common involvement of mitochondrial dysfunction in HLHS regardless of HF outcomes. This is supported by another study that also reported mitochondrial defects in HLHS iPSC-CM (Paige *et al*., 2020). With the outcome-based iPSC-modeling, we showed the mitochondrial dysfunction and oxidative stress underlie the early HF in HLHS, while a hyper-elevated antioxidant response may provide protection from oxidative and ER stress to prevent early HF. Together these findings suggest early HF is the result of uncompensated mitochondrial mediated oxidative stress. The observed altered regulation of YAP1 suggests the tantalizing possibility that the mitochondrial defects also may contribute to the LV hypoplasia in HLHS, a question that warrants further studies.

We also showed possible therapeutic intervention with the targeting of mPTP closure with sildenafil or suppression of UPR with TUDCA. We note Sildenafil is already being used empirically to threat HF associated with pulmonary hypertension(Guglin et al., 2016). Suppression of UPR, such as with TUDCA, may be another therapeutic path. TUDCA is currently in clinical trial for amyotrophic lateral sclerosis(Elia et al., 2016). Providing antioxidant might be another therapeutic course, although we found ascorbic acid did not rescue mitochondrial defects in the Group II iPSC-CM. Overall, our iPSC modeling has yielded new insights into the underlying causes for early HF in HLHS and suggest new evidence-based therapies that will need to be further investigated. These findings suggest a new paradigm for modeling clinical outcome using patient stratified iPSC.

### Limitations of the Study

One limitation of our study is the inclusion of iPSC-CM from only 10 patients. However, this compares favorably to other studies that typically include iPSC from only one to three patients, and no study had controlled for outcome(Gaber *et al*., 2013; Hrstka *et al*., 2017; Jiang *et al*., 2014a; Kobayashi *et al*., 2014; Miao *et al*., 2020; Paige *et al*., 2020). Nevertheless, the generalizability of our findings will require future confirmation with analysis of iPSC-CM from additional patients. As our study was focused on acute early-HF in patients less than one year old, the relevance of these findings to HF in older HLHS patients will require further studies. While additional factors may contribute to HF in older patients, the involvement of mitochondrial dysfunction is likely. This is suggested by the recovery of mitochondrial-related pathogenic variants in the expanded WES analysis of 41 HLHS patients that included older patients with heart transplant.

## Supporting information

all main text

## ACKNOWLEDGMENTS

This work was supported by funding from University of Pittsburgh (C.W.L), NIH HL132024 (CWL), HL142788 (CWL, MT), NIH HL144776 (GP), DOD PR140183 (CWL), postdoctoral fellowship (X.X.) jointly funded by the American Heart Association and the Children’s Heart Foundation. Some human heart tissues were obtained from the Molecular Atlas of Lung Development Program (LungMAP) Consortium distributed by Human Tissue Core (HTC) supported by NIH grants HL122700 and HL148861 (G. H. Deutsch, T. J. Mariani, and G. S. Pryhuber). Donor tissue was supplied through the United Network for Organ Sharing. We thank the staff of the HTC including Heidie Huyck and Cory Poole. We are grateful to families who generously give such precious gifts to support this research.

## AUTHOR CONTRIBUTIONS

Study design: C.W.L. and X.X; miPSC and hiPSC reprogramming: iPSC-CM differentiation, cardiomyocyte proliferation and apoptosis, cardiomyocyte and mitochondrial function measurements, drug screening and data analysis: X.X; Single cell RNAseq and data analysis: X.X,K.J,B.A,H.Y.,C.W.L.,A.S.B,D.K; human patient clinical data analysis: J.I.L,P.A; recruitment of subjects and human tissue sample collection: C.W.L.,J.I.L.,P.A.,G.B, G.A.P; human heart tissue analysis: G.B,G.A.P.,X.X.; Seahorse measurement support: S.S.S; mouse fetal ultrasound imaging and mouse phenotyping: X.L.,X.X.; mitochondrial staining support and analysis: X.X.,T.N.F.G.A.P; iPSC-CM sarcomere video analysis: P.N, J.C, C.K.K; human exome sequencing analysis: W.Z; protein network analysis, K.B.K, M.K.G.; statistics: X.X; manuscript preparation: C.W.L, X.X, A.S.B, K.J, B.A, G.A.P., J.I.L, R.A.D, M.T., M.K.G., W.Z and T.N.F.

## EXPERIMENTAL MODEL AND SUBJECT DETAILS

### Mouse Strain

E13.5-E14.5 Ohia HLHS mouse (*Sap130m/m;Pcdha9m/m*) or CRISPR HLHS mouse (*Sap130m/m;Pcdha9m/m*) and littermate controls were used for primary cardiomyocytes explants from heart tissue. Mouse embryo fibroblasts used for mouse iPSC generation were generated from E14.5 – 17.5 mouse embryos (**See Supplemental Spreadsheet1**). All mice were housed, treated, and handled in accordance with the guidelines set forth by the University of Pittsburgh Institutional Animal Care and Use Committee and the National Institutes of Health’s Guide for the Care and Use of Laboratory Animals.

### Human Blood, Cells, and Surgical Tissue

Cells, heart tissue and blood were obtained from HLHS patients recruited from Children’s Hospital of Pittsburgh of UPMC with informed consent under a human study protocol approved by the University of Pittsburgh Institutional Review Board (Supplemental Spreadsheet 1). For infants and minors, informed consent was obtained from the legal guardian. Some human heart tissues were obtained from the Molecular Atlas of Lung Development Program (LungMAP) Consortium distributed by Human Tissue Core (HTC). Donor tissue was supplied through the United Network for Organ Sharing for Western blot and isolated mitochondrial OCR measurements.

## METHOD DETAILS

### Production of patient iPS cells

Mouse embryonic fibroblasts were reprogrammed using the CytoTune-iPS Sendai Reprogramming kit(Fusaki et al., 2009). Human fibroblasts or lymphoblastoid cells were transfected with four episomal plasmids(Okita et al., 2011) using electroporation. iPSCs clones were identified by immunofluorescent staining of pluripotency marker Oct4 and Nanog and qPCR analysis of stem cell markers **(Supplemental Figure S2B)**(Xu et al., 2013). All antibody and primer sequence information are provided in **Supplemental Spreadsheet 1**

Several independent iPSC clones were isolated for four of the HLHS patients, and analysis conducted with these independent clones generally yielded similar results (see **Figure S2D**). While independent iPSC clones from one subject are often used to demonstrate reproducibility of findings, one study using transcriptome profiling showed the importance of using iPSCs of different parental origin rather than multiple sister iPSC clones to distinguish disease-associated mechanisms from genetic background effects in disease modeling(Schuster et al., 2015).

### Production of iPS derived cardiomyocytes

The iPSC cells were seeded on BD Matrigel pre-coated plates for 2-3 days under mTESR1 media then switched to CDM3 media consisting of RPMI 1640, BSA, 213 μg/ml Vitamin C (Ascorbic acid) and 6 μM CHIR99021(Burridge et al., 2014). After 2 days the media was replaced with CDM3 Media containing RPMI 1640, BSA, 10 μM XAV939, 213 μg/lm Vitamin C, and BSA. Finally, ∼14 days after initiating reprograming, beating cells are observed and further analyzed in the following days (Day18-22).

### Immunofluorescence Staining

Cells were fixed with 4% paraformaldehyde with 0.1% Triton X-10, followed by blocking in 5% goat serum, then staining overnight with primary antibody in 0.5% bovine serum albumin/phosphate-buffered saline (BSA/PBS). After washing in PBS, incubation with secondary antibody was performed in 0.5% BSA/PBS for 1 hour at room temperature and nuclei were stained with 2 μg/ml Hoechst 33342 (Life Technologies). Images were acquired using the Leica SP8 confocal or Leica DMI6000-SD microscopes. Antibody information in **Supplemental**

### Spreadsheet 1

#### Analysis of mitochondrial calcium transients

iPSC-CMs cultured in chamber slides were loaded with 1 μM Rhod-2 (Molecular Probes, Life Technologies, Carlsbad, CA, USA) in Hank’s balanced salts modified buffer (HBSS, pH 7.4) for 15 minutes at 37°C and washed twice for 15 minutes in HBSS. The slides were placed on a temperature-regulated microscope stage and kept at 37°C. Fluorescence images were acquired using the ImageJ time series analyzer package (NIH, Bethesda, MD; Version: 2.0.0-rc-69/1.52K) together with Leica DMI6000-SD fluorescence microscope. The data shown represent the average of Rhod-2 intensity for 3 controls and 10 HLHS patients iPS-CMs from three independent experiments.

### Analysis of sarcomere contractility in iPSC-CM

Single iPSC-CM cell videos were collected by Leica DMI 3000B microscope and videos of human iPSC-CM (200 Hz) containing striated sarcomere were analyzed using a custom MATLAB code (available upon request) written to apply the fast Fourier transform (FFT) algorithm to each frame (approximately 1400 frames per video) to compute the spatial frequencies of the sarcomeres. The frequency (f_0_) of the highest amplitude peak of the FFT within a user defined range was identified, and sarcomere length (L) in each frame was calculated by taking the reciprocal of f_0_, 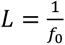. The user defined range was determined and optimized to ensure that the maximum and minimum measured sarcomere lengths always occurred within this range. The sarcomere length in units of pixels was then converted to units of micrometers, 1µm = 4.58 pixels (100X magnification video), and plotted as a function of time (seconds). From the plots of sarcomere length versus time, the point of maximum sarcomere length immediately before contraction (t_1_, max), minimum sarcomere length (Systolic length) during contraction (t_2_, min), and maximum sarcomere length (Diastolic length) immediately after contraction (t_3_, max) were identified for each contraction. Fractional shortening (FS) was calculated as 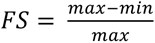, contraction rate (CR) was calculated as 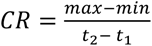, and relaxation rate (RR) was calculated as 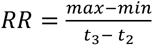.

### RNA extraction, real-time PCR, and transcript splicing analysis

Total RNA was isolated using the miRNeasy micro-Kit (QIAGEN) with on-column DNase I digestion (QIAGEN). cDNA was prepared with high-capacity RNA to cDNA kit (Applied Biosystems). Real-time PCR was conducted using 7900HT Fast Real Time PCR System. All primer sequence information is provided in **Supplemental Spreadsheet 1**

### Seahorse Analyzer analysis of oxygen consumption rate

For cell oxygen consumption rate (OCR) quantification, 20,000 iPS-CMs or 20000 iPSCs were seeded into each well of a Seahorse XFe96 cell culture plate and cultured for 2 days for adherence to the culture plate. On day of measurement, the medium was changed to pre-warmed Seahorse assay medium, and OCR determined using the Seahorse XF Cell Mito Stress Kit (Agilent). Basal respiration was measured in unstimulated cells. Afterwards, oligomycin (1 μM) was added to quantify respiration coupled to ATP production and proton leak followed by carbonyl cyanide-4-(trifluoromethoxy)-phenylhydrazone (FCCP; 1 μM) injection to assess maximal cellular respiration (respiratory capacity). Finally, antimycin A (1 μM) and rotenone (1 μM) were used to assess non-mitochondrial respiration. For mouse and human heart tissue OCR quantification, 2mm X 2mm heart tissue pieces were seeded into each well of a Seahorse XF24 islet capture microplate and OCR were measured using the same Seahorse XF Cell Mito Stress Kit to obtain the basal respiration rate in unstimulated cells during two cycles of measurement.

### Analysis of inner mitochondrial transmembrane potential

Embryonic left and right ventricles were dissociated with papain to generate primary cardiomyocytes for live imaging as previously described (Hom *et al*., 2011). Briefly, this entailed loading live explanted cardiomyocytes, iPSC, and iPSC-CM for 35 minutes with tetramethylrhodamine ethyl ester (TMRE, 20 nM, Invitrogen, Cat# T-669) and Mito Tracker Green (MTG, 200 nM, Invitrogen, Cat# M-7514) in Hepes-Tyrode’s buffer, washed, and equilibrated for 20 minutes in the same buffer. The live cells were then imaged using epifluorescence microscopy. Mitochondrial membrane potential (Δψ_m_) was quantified as the ratio of TMRE to MTG intensity (Galmiche et al., 2011).

### Mitochondrial network analysis

MitoTracker Red (100 μM) was loaded into live cells following manufacturer’s recommendations, and then cells were fixed in 4% paraformaldehyde/PBS at 37°C for 15 minutes. Cells were permeabilized in 0.2% Triton X-100/PBS for 10 minutes and then immunostained for cTnT while DNA was labeled with Hoechst. Mitochondria were imaged using a Leica SP8 confocal with a 40x / 1.3NA objective. Acquisition settings and deconvolution were done with the guidance of SVI Huygens software, and images were post-processed in ImageJ (NIH, Bethesda, MD; Version: 2.0.0-rc-69/1.52K) with unsharp mask (radius 2; mask weight 0.7), background subtraction, and the tubeness filter (sigma = 0.25 microns) to highlight mitochondrial filaments. Mitochondria were segmented with Skeletonize 2D/3D. Mitochondrial networks were then analyzed (“Analyze Skeleton”) using BoneJ. Clusters with 20 branches or more were used for measuring average branch length and linkage statistics. To quantify the degree of mitochondrial consolidation, clusters were ranked from most to least branches (see graphs) and a mono-exponential decay curve is fit to the resulting data. The curve’s decay constant is then inverted so that higher values reflect more-linked mitochondrial networks.

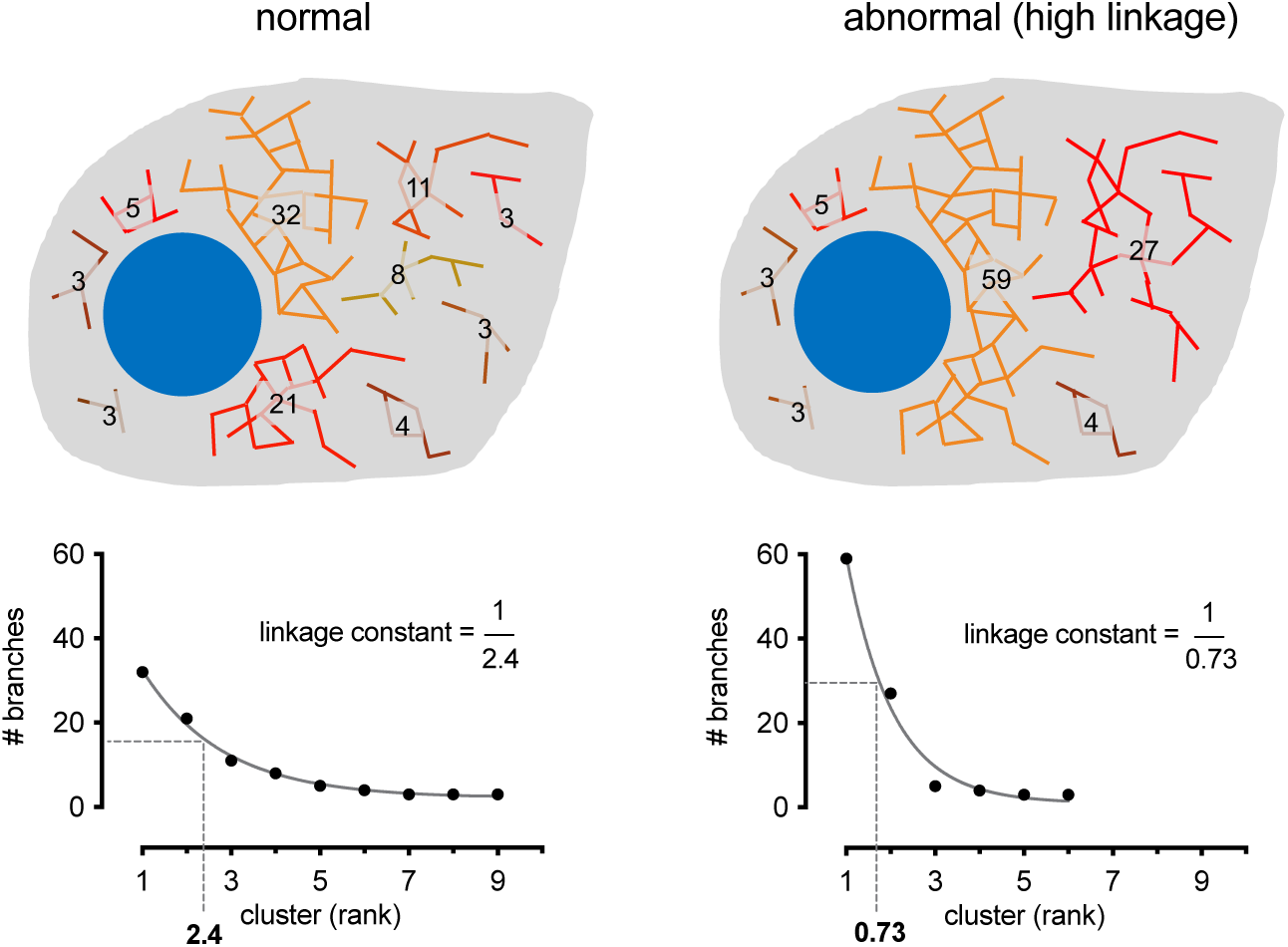

### Mitochondrial DNA copy number assays

DNA was extracted from human iPSC-CM. qPCR was performed and mitochondrial DNA copy number was determined by normalizing results from primers targeted to mtDNA-tRNA-Leu (Forward: 5’-CAC CCA AGA ACA GGG TTT GT-3’ and Reverse: 5′-TGGCCATGG GTA TGTTGT TA -3′) against results from primers targeted to nuclear B2-microglobulin (Forward: 5’-TGCTGT CTC CAT GTT TGA TGT ATC T-3’ and Reverse: TCT CTG CTC CCC ACC TCT AAG T-3’(Rooney et al., 2015).

### Reactive oxygen species, nitric oxide and lysosome measurements

To quantify reactive oxygen species (ROS), nitric oxide level and lysosome abundance, iPSC-CMs were incubated at 37 °C for 30 minutes with 5 μM MitoSOX, 5 μM DAF-FM diacetate and 1 μM LysoTracker Red DND-99 (Life Technologies), respectively. For NO measurement, additional 15–30 minutes incubation could complete de-esterification of the intracellular diacetates. CD172a(SIRPα/β) was used as a human iPSC-CM marker (Dubois et al., 2011). Live cell fluorescent imaging was conducted using the Leica DMI6000-SD microscope.

### Human heart tissue and iPSC-CM Western blotting

LV and RV tissue or iPSC-CM were homogenized and processed for Western blotting using a ChemiDoc (Biorad) with Image J image processing (Beutner et al., 2017; Beutner et al., 2014). Antibodies from Abcam and BioRad were used and included: OXPHOS Rodent Cocktail (ab110413), AC (#154856), Starbright 700 (anti-mouse), Starbright 520 (anti-rabbit).

### Isolation of mitochondria and oxygen consumption assay

Mitochondria where isolated on ice from fresh or frozen tissue (∼140 mg) in isolation medium by homogenization and differential centrifugation and resuspended in EGTA/EDTA-free isolation (Beutner *et al*., 2017; Beutner *et al*., 2014). Oxygen consumption was measured at room temperature in respiration medium with a Clark oxygen electrode (Hansatech) using published protocols. Cytochrome c (50µM) and atractyloside (100µM) were used to test mitochondrial membrane integrity. Substrate-mediated respiration (state 2 or V0), maximal respiration (state 3 or Vmax), and RCR (Vmax over V0) were calculated.

### Single-cell RNA sequencing

Previous study proved there is no significance difference between iPSC-CM from day 21 and day 30(funakoshi et al., 2018), hence, the iPSC-CM differentiated at day 22 were choose as scRNAseq samples. The iPSC-CM from three patients and one control was prepared for single cell RNAseq. The iPSC-CMs were disaggregated using cold active protease [10 mg/ml Bacillus Licheniformis protease; Creative Enzymes NATE0633) and 125 U/ml DNase (Applichem, A3778) incubated on ice with trituration 5-7 minutes, then 5% bovine serum albumin (BSA) was added, and cells were filtered by 100 μm cell strainer and the cells pelleted, then re-suspended in 200ul PBS/BSA. Trypan blue exclusion was used to quantify cell viability, and the volume was adjusted to 200,000 cells/ml for 10X chromium single-cell RNA-seq. Pair-end library preparation was carried out using the V3 version (10X Genomics). Single-cell droplet libraries from ∼10K cells from each suspension were generated using the 10X Genomics Chromium controller with the Chromium Single Cell 3’ GEM Library and Gel Bead Kit v.3 and the Chromium Chip B Single Cell kit (1 GEMs per sample, expected recovery ∼6k cells per GEM). All samples were barcoded with the Chromium i7 Multiplex Kit. All libraries were pooled and sequenced across two lanes of a HiSeq4000, 150bp paired end reads with a target coverage of 20k fragments per cell. All samples were uniquely indexed, mixed, and evenly distributed into the Illumina HiSeq 4000 for sequencing.

### Single-cell RNA-Sequencing Data Analysis

Single-cell sequencing data was processed using the Cell Ranger (version 3.1.0) count pipeline using the human reference genome GRCh38 and annotations from Ensembl (version 93). Quality control and filtering were performed using scater (McCarthy et al., 2017) (v1.18.6). For each sample, cells with library size less than 500, number of detected genes less than 300 or greater than 6,000, or mitochondrial percentage greater than 4 times the median absolute deviation (MAD) from the median value were excluded. Additionally, top 3% cells ranked by the doublet score (hybrid) calculated using the scds R package (Bais and Kostka, 2020) (v1.6.9) were excluded. Only non-ribosomal genes with at least 1 count in ≥ 5 cells were considered. We adapted the approach of Kannan et al. [https://doi.org/10.1101/2020.04.02.022632] for cell type classification using SingleCellNet(Tan and Cahan, 2019) (v0.1.0) and further limited to cells classified as “cardiac muscle cells” yielding a data for 13,954 genes across 8,094 cells. We performed downstream analyses using the Seurat package (Stuart *et al*., 2019). To focus on high-quality CMs, we further removed cells with total library size less than 1,400 or number of detected genes ≤800, or percentage of mitochondrial gene counts greater than 20%. This yielded a final set of 877, 1,718, 1,434, 374 cells for 1053, 7042, 7052 and 7464, respectively, for downstream analysis.

We normalized total count per cell to 10,000 and find top 2000 highly variable genes in each sample. Integration of cells from different samples and batch correction were performed using IntegrateData function in Standard procedure of Seurat 3. Scaled data after integration was used for principal component analysis (PCA) and top 30 dimensions were used for neighbor detection and Louvain clustering (resolution = 0.5). UMAP was drawn for the visualization of single-cell data in reduced dimensions.

Differentially expression analysis was conducted using student t-test in Scanpy(Wolf et al., 2018). We compared differentially expressed genes of clusters and sample groups, as well as samples and sample groups per cluster (**Figure 5C-F,G**, S5F). Genes with FDR < 0.05 in tests were selected as DEGs. ToppGene(Chen et al., 2009) was used for gene enrichment analysis and Gene Ontology (Biological Process) terms and coexpression of MSigDB were used for annotations of gene lists. The strength of associations was represented by -log10(FDRToppGene) (**Figure 5C-F,G**). Gene modules of cardiomyocyte clusters were generated using 200 most significantly upregulated genes (Figure S5E) and their top enriched Gene Ontology (Biological Process) terms were used for annotating cluster identities. Similarity between these clusters were evaluated using Pearson correlation of genes in gene modules. Cell cycle scores were calculated in Seurat using CellCycleScoring function and cell cycle phases were inferred accordingly (Figure S5C).

### Whole exome sequencing analysis

Whole-exome capture was carried out on 6 Caucasian HLHS subjects with iPSC and 41 HLHS subjects (including the 6 HLHS subjects) at BGI Americas. Genomic DNA from venous blood was captured with Agilent V4 Exome Capture kit. Sequencing was performed on the Illumina HiSeq2000 platform with 100 paired-end reads, or the Illumina HiSeq4000 with 150 paired-end reads at 100× coverage. Sequence reads were mapped to the reference genome (hg19) with BWA-MEM(Arakawa et al., 2010) and further processed using the GATK(McKenna et al., 2010) Best Practices workflows, which include duplication marking, and base quality recalibration. Single nucleotide variants (SNVs) and small indels (InDels) were detected using GATK haplotypeCaller and annotated by Annovar(Wang et al., 2010). High quality variants were recovered that: 1) passed GATK Variant Score Quality Recalibration (VSQR); 2) have minimum 5 supported reads; 3) have genotype quality ≥ 20 or 60 for SNVs or InDels, respectively; 4) SNVs or InDels not within 10bp or 5bp of an indel, respectively.

Variants with minor allele frequency (MAF) was less than 0.01 in GnomAD exome (Karczewski et al., 2020)or Kaviar database were retained for downstream analyses. Only loss-of-function (LoF) mutations (nonsense, canonical splice-site, frameshift indels, and start loss), likely damaging missense variants (D-Mis) and non-frameshift indels were considered potentially damaging. Missense variants were considered likely damaging if it was predicted to be damaging by at least three out of nine prediction scores available via dbNSFP v3.5a (Liu et al., 2016). All filter processes are shown in **Figure S7.**

### Functional enrichment and interactomes analysis

Webgestalt KEGG pathway analysis (http://webgestalt.org/) was performed for unique LoF variants from 6 HLHS cohort. The interactomes of the four genes harboring unique variants in the HLHS unfavorable patient was assembled by including their protein-protein interactions (PPIs) collected from BioGRID (Stark et al., 2011) and HPRD (Prasad et al., 2009), and novel PPIs predicted by High-precision PPI Prediction (HiPPIP) model (Ganapathiraju et al., 2016) focusing on short path connections to the Hippo pathway. Hippo pathway genes were extracted from KEGG (Kanehisa et al., 2008). Enrichment Analysis Tool available on Gene Ontology (GO) website which uses PANTHER (Ahmad et al., 2013) on the backend, was used to find biological process terms associated with the interactome genes with statistical significance. It computes fold enrichment of the genes in the input list over the expected value. Fold enrichment>1 and fold enrichment<1 showed that the annotation is overrepresented and underrepresented in the list respectively. It presents p-value determined by Fisher’s exact test with FDR correction, and a cut-off of 0.05 was used to select significantly enriched annotations.

## QUANTIFICATION AND STATISTICAL ANALYSIS

Standard statistical analyses were performed using GraphPad Prism 9. D’Agostino & Pearson normality test and Shapiro-Wilk normality test were used to test if the data had a Gaussian distribution. For Gaussian distribution, data are presented as bar graphs and expressed as mean±SEM, either Unpaired t-test (Two-tailed) or One-way ANOVA (FDR B&Y correction were used for multiple comparisons) were applied. Data without Gaussian distribution showed by box plot (Line at median and minimum-maximum were represented by the top /bottom of box), either non-parametric Mann-Whitney test (Two-tailed) or Kruskal-Wallis tests (FDR B&Y correction were used for multiple comparisons) were used. The experiments were not randomized. The investigators were not blinded to allocation during experiments and outcome assessment.

## SUPPLEMENTAL INFORMATION

**Supplemental Figure 1: Mitochondrial defects in the HLHS mouse heart tissue and HLHS mouse iPSC-CM**. Related to Figure 1.

**Supplemental Figure 2: Generating iPSC and iPSC-CM from HLHS patients and control subjects Related to Figure 1.**

**Supplemental Figure 3: Mitochondrial defects and altered Hippo signaling Related to Figure 2&3.**

**Supplemental Figure 4: Inhibition of the mitochondrial permeability transition pore rescues mitochondrial respiration and YAP1 nuclear localization. Related to Figure 4.**

**Supplemental Figure 5: Analysis of HLHS patient iPSC-CM using single-cell RNAseq Related to Figure 5.**

**Supplemental Figure 6: Protein-Protein Interactome of Genes Highly Expressed Only in Patient 7052. Related to Figure 5.**

**Supplemental Figure 7: Whole exome sequencing analysis of pathogenic variants show enrichment for metabolic-mitochondrial pathways in Group II HLHS patients Related to Figure 7.**

**Supplemental Spreadsheet 1: Patient information, iPSC production, antibodies, and primer sequences. Related to STAR Methods.**

1. miPSCs generated and used in this study
2. HLHS patient medical history
3. Patient iPSCs reprogramming
4. Human and mouse primer sequences.

**Supplemental Spreadsheet 2: Single cell RNAseq related information. Related to Figure5,6.**

1. Marker gene list for C0-8 (Figure S5 E)

2.1-2.9. GO enrichment analysis for Cluster 0-8 (Figure S5 E)

2.10. HCM related genes in Clusters C3 and C8 (Figure 5 A)

3.1. DEG No. in Each Cluster (Figure 5 B)

3.2-3.10. GO enrichment analysis of DEG in C0/C1/C5 under different comparisons (Figure 5 C-E)

4.1 All DEGs in Figure 5F

4.2 Toppgene of Control 1053 DEGs (Figure 5F)

4.3 Toppgene of Patient 7464 DEG (Figure 5F)

4.4 Toppgene of Group II 7042/7052 Shared DEG (Figure 5F)

4.5 Toppgene of Patient 7042 DEG (Figure 5F)

4.6 Toppgene of Patient 7052 DEG (Figure 5F)

5.1. Mitochondrial DEGs (Figure 5H)

5.2 Toppgene of Control 1053 Mitochondrial DEGs (Figure 5H)

5.3. Toppgene analysis of Patient 7464 Mitochondrial – DEGs (Figure 5H)

5.4. Toppgene of Group II Mitochondrial-DEG (Figure 5H)

5.5. Toppgene of Patient 7042 Mitochondrial-DEG (Figure 5H)

5.6. Toppgene of Patient 7052 Mitochondrial-DEG (Figure 5H)

6. Patient 7052 - 28 upregulated DEGs (Figure 5G)

7.1. Protein-Protein Interactome network genes (Figure S6A)

7.2. PPI BiNGO Biological Process Pathway Enrichment (Figure 5I)

8.1-8.2. GO enrichment analysis of DEG of Group II VS Control in C7 (Figure 6A)

**Supplemental Spreadsheet 3: Whole exome sequencing and interactome analysis related information. Related to Figure7.**

1. Description
2. Unique LoF genes from Group II patient (Figure 7A)
3. LoF (loss of function) variants from Group II patient (Figure 7A)
4. Webgestalt/KEGG pathway enrichment of unique LoF genes from Group II patient (*OXA1L, NNMT, NEU3, ALDH7A1*) (Figure 7A)
5. Protein-protein interactome GO Biological Processes
6. GO Biological Processes in Figure 7B
7. Genes in GO Biological Processes in Figure 7B.
8. Unique gene with variants in 41 HLHS cohort (Figure 7C)
9. Extreme variant list in 41 HLHS cohort (Figure 7C)
10. Metascape-GoEnriched (Figure 7C)
11. Mitochondrial gene list (Figure 7D)
12. Damaging variants in mitochondrial-related genes (Figure 7D)
13. Overlapping mitochondrial and unique genes in 41 HLHS cohort (Figure 7D)
14. Toppgene analysis of those 19 overlapping genes (Figure 7D)

## SUPPLEMENTAL VIDEO LEGEND

**Supplemental Videos**

Sup-video-1_hips-cm_beating-Related to Figure 1

Sup-video-2_hips-m_Ca-Related to Figure 1

Sup-video-3_hips-cm_single_cell-Related to Figure 1

**Sup-video-1_hips-cm_beating:** Videomicroscopy showing contraction of human iPSC-CM. The iPSC-CM from control subject and Group I beat faster than iPSC-CM from Group II. Scale bar = 250 μm.

**Sup-video-2_hips-cm_Ca:** Calcium transients in the iPSC-CM are visualized using Rhod-2. Note faster propagation of calcium transients in iPSC-CM from control subject and Group I patients as compared to that of Group II. Scale bar = 250 μm.

**Sup-video-3_hips-cm_single_cell:** Videomicroscopy recording of individual beating iPSC-CM from control subject, Group I and Group II patients. Robust contractions are seen in cardiomyocytes from control and Group I, but only weak contractions are seen in Group II. Scale bar = 10 μm.

