## Supplementary material for "iPSC modeling shows uncompensated mitochondrial mediated oxidative stress underlies early heart failure in hypoplastic left heart syndrome": all main text

### SUPPLEMENTAL FIGURE LEGEND

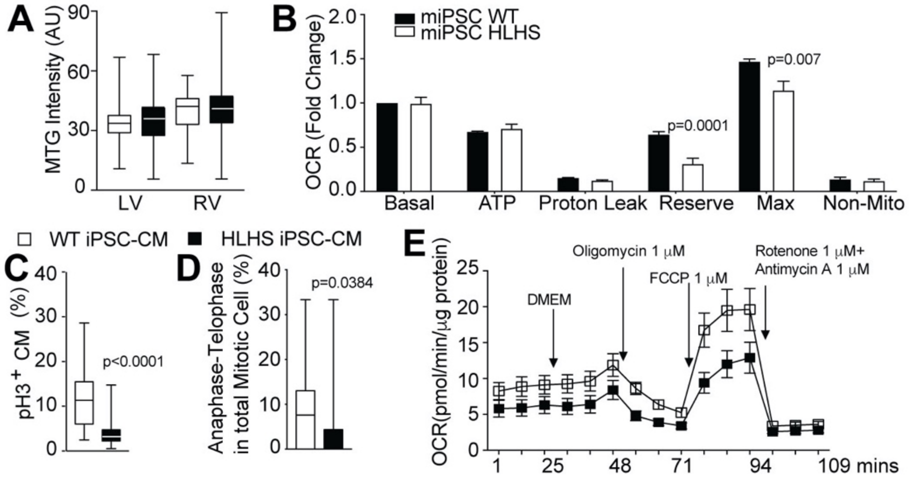


**Figure S1. Mitochondrial defects in the HLHS mouse heart tissue and HLHS mouse iPSC-CM.** Related to Figure 1.

(A) Mitotracker green (MTG) was used to quantify total mitochondrial mass in primary cardiomyocyte explants from the LV and RV of *Ohia* HLHS mutant and littermate wildtype fetuses. No significant difference was observed between the HLHS mutant vs. wildtype LV or RV.

(B) Oxygen consumption rate (OCR) measured using the Seahorse Analyzer showed differences between wildtype vs. HLHS mouse iPSC in respiratory reserve and maximal OCR.

(C) Quantitative analysis of pH3 expression showed reduction in proliferation of the *Ohia* mouse HLHS iPSC-CM.

(D) Proportion of mouse iPSC-CM cells in anaphase/telophase as a percentage of total iPSC-CM in mitosis was reduced in the HLHS iPSC-CM.

(E) Profile of oxygen consumption related to oxidative phosphorylation as observed with the Seahorse Analyzer in wildtype and *Ohia* HLHS mouse iPSC-CM. This entails conducting a metabolic stress test as follows. First baseline OCR measurement is obtained, followed by the addition of 1 mM oligomycin and three OCR measurements, then the injection of 1 mM FCCP followed by another three OCR measurements, and finally 1 mM rotenone and 1 mM antimycin A addition and three more OCR measurements. FCCP, carbonyl cyanide 4-(trifluoromethoxy)phenylhydrazone.

Bar graphs show mean±SEM with Student's t-test. Box plots show median and minimum-maximum, with Mann-Whitney statistical test. (A) WT-LV (n=124 CMs) and RV (n=86 CMs), *Ohia* HLHS mutant-LV (n=105 CMs) and RV (n=118 CMs) (WT=3 and HLHS=3 litters). (C, D) WT n=2800 CMs and HLHS n=1800 CMs. (B,E) WT/HLHS n=2 lines and 3 independent repeats.

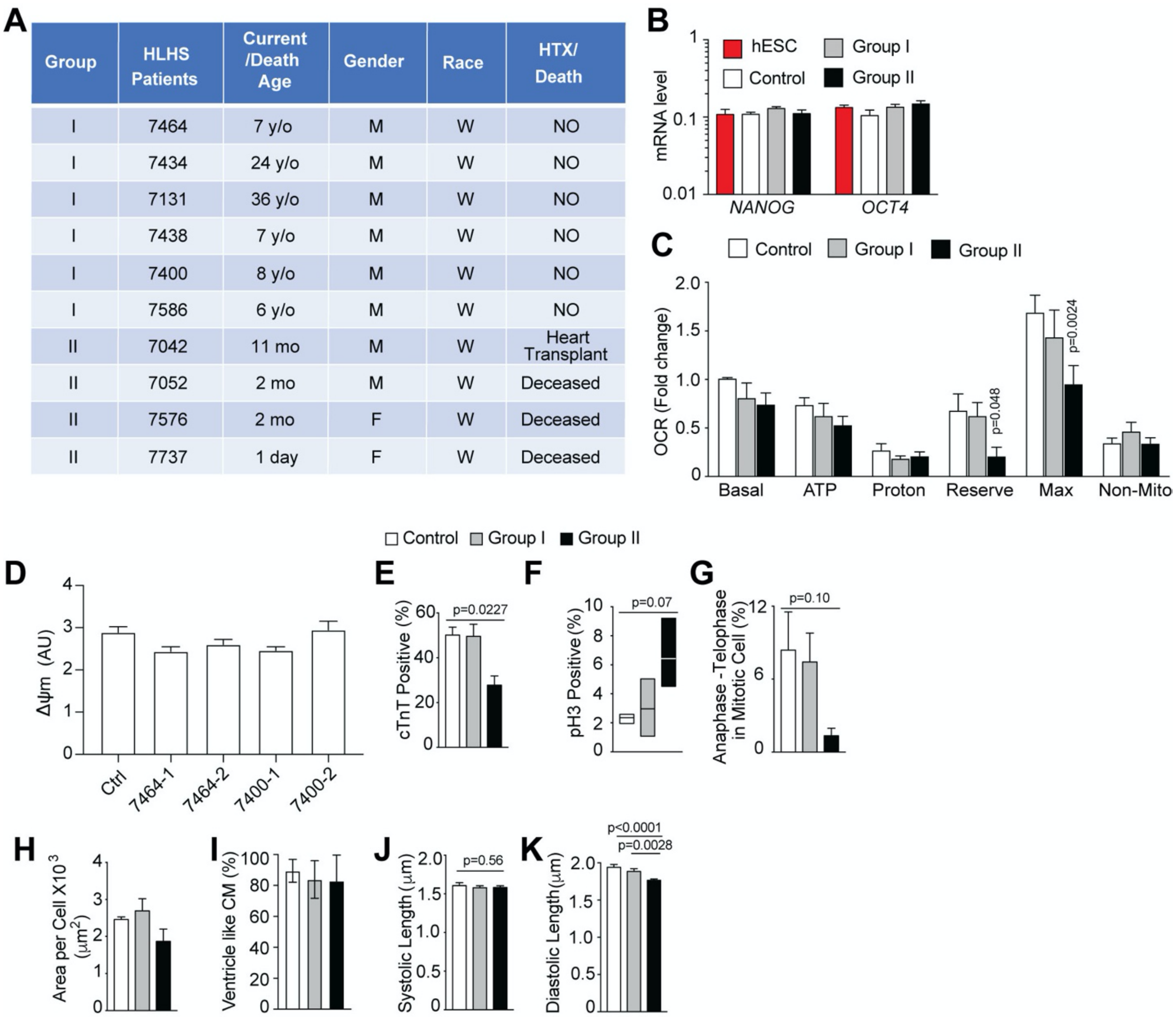


| Group | HLHS Patients | Current /Death Age | Gender | Race | HTX/ Death |
|---|---|---|---|---|---|
| I | 7464 | 7 y/o | M | W | NO |
| I | 7434 | 24 y/o | M | W | NO |
| I | 7131 | 36 y/o | M | W | NO |
| I | 7438 | 7 y/o | M | W | NO |
| I | 7400 | 8 y/o | M | W | NO |
| I | 7586 | 6 y/o | M | W | NO |
| II | 7042 | 11 mo | M | W | Heart Transplant |
| II | 7052 | 2 mo | M | W | Deceased |
| II | 7576 | 2 mo | F | W | Deceased |
| II | 7737 | 1 day | F | W | Deceased |

**Figure S2. Generating iPSC and iPSC-CM from HLHS patients and control subjects** Related to Figure 1.

(A) Basic demographic information on the ten HLHS patients selected for the generation of iPSC and their clinical outcome with death or heart transplantation (HTX) indicated.

(B) qPCR showed similar *NANOG* and *OCT4* transcript levels in iPSC from Group I and II HLHS patients vs. control subject iPSC or human embryonic stem cells (hESC).

(C) Assessment of mitochondrial respiration parameters in control, Group I and Group II undifferentiated iPSC showed little or no change except for respiratory reserve and maximal OCR for Group II iPSC.

(D) Mitochondrial membrane potential was measured in different sister iPSC lines from the same patient.

(E) Reduction in cTnT positive cells in Group II but not Group I iPSC-CM.

(F,G) Group II iPSC-CM showed increased mitosis (F) accompanied by a marked decrease in the proportion of mitotic cells in anaphase/telophase (G), indicating a block in mitotic cell cycle progression.

(H) Analysis of cell size in the immunostained cTNT positive iPSC-CM showed no difference in cell size between the iPSC-CM of HLHS patients vs. the control subjects.

(I) Analysis of the calcium transient trace to identify atrial vs. ventricular myocytes showed.the majority of iPSC-CM are of ventricle phenotype.

(J) Systolic (Minimum) and (K) diastolic (Maximum) sarcomere length was quantified with analysis of video microscopy recording of contractile motion in individual iPSC-CM.

Bar graphs show mean ± SEM with one-way ANOVA. (B,C) n=3,6,4 subjects. (D) n=57, 43, 85, 118, 47CMs. (E,G,H) n=3,5,3 subjects. (F) n=3,6,3 subjects. (I) n=3,6,4 subjects. (J,K) n=17, 23, 38 single cells from 3 controls, 4 group I and 4 group II subjects.

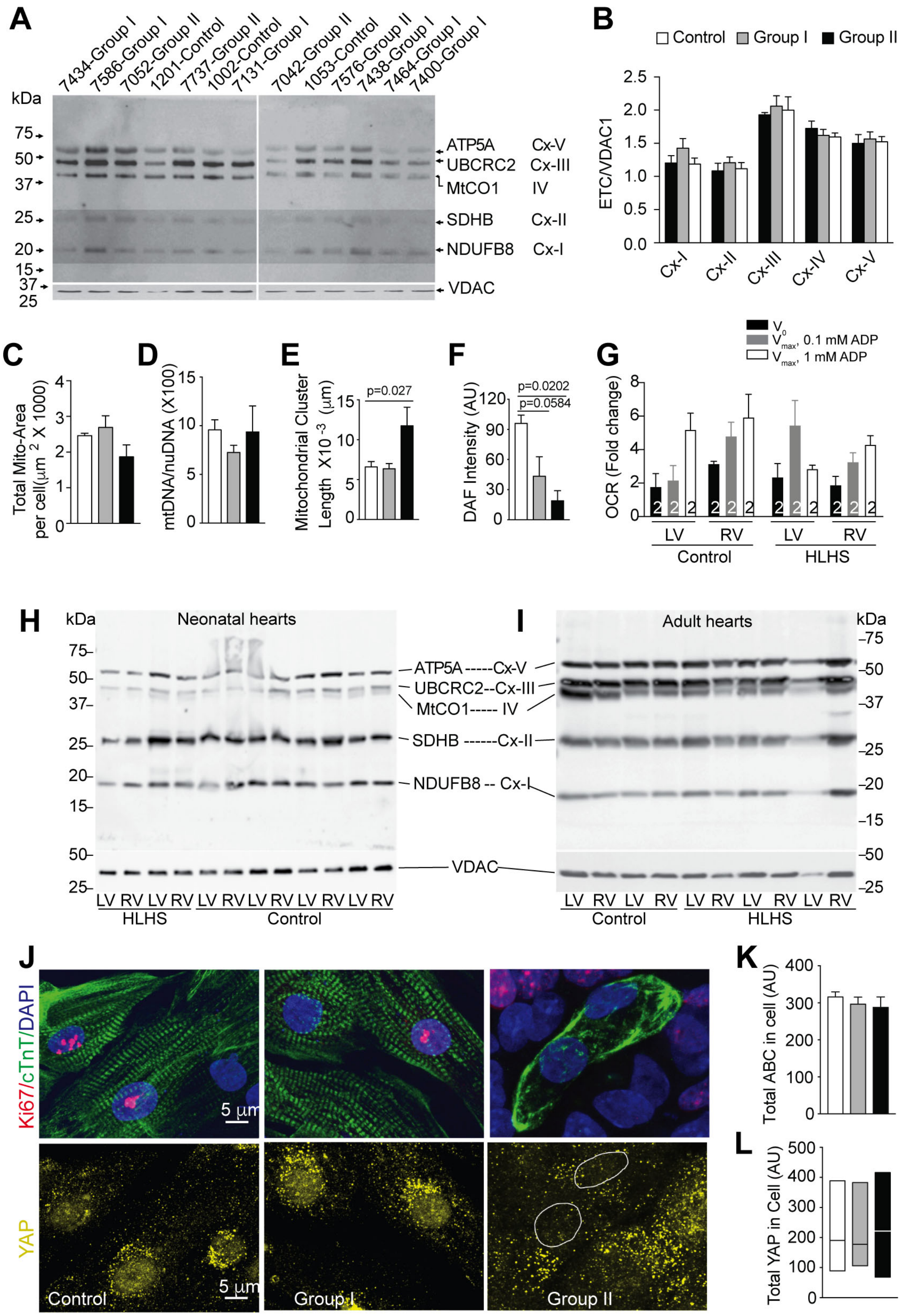


**Figure S3. Mitochondrial defects and altered Hippo signaling** Related to Figure 2&3.

(A, B) Immunoblots (A) and quantification (B) of electron transport chain (ETC) proteins and voltage dependent anion channel (VDAC) from control subjects, and Group I and Group II HLHS patient iPSC-CM. Quantification of the Western blots showed no significant difference between the different groups (statistical analysis by ANOVA). Note: an increased contrast was used to enhance visualization of SDHB and NDUFB8 in panel (A).

(C) Mitotracker Red staining of mitochondria showed no change in total mitochondrial area between control and Group I and II iPSC-CM.

(D) Measurement of mitochondrial DNA/nuclear DNA ratio by qPCR showed no difference between control, HLHS Group I and Group II iPSC-CM.

(E) Confocal imaging showed average length of mitochondrial cluster was increased in the iPSC-CM of the HLHS Group II but not Group I as compared to control.

(F) DAF staining intensity was used to quantify NO levels in iPSC-CM. This showed significant decrease in NO in the Group II iPSC-CM, and with a trend for lower NO in the Group I iPSC-CM.

(G) Maximal oxygen consumption (Vmax) after addition of succinate (V0) followed 0.1 mM or 1 mM ADP in isolated mitochondria extracted from 2 explanted human neonatal control and 2 HLHS patient hearts.

(H,I) Immunoblots for ETC proteins and VDAC from human LV and RV heart tissue. Panel (H) show results from neonatal control (N=4) and HLHS (N=2) hearts, and panel (I) shows adult control heart (N=2) and HLHS heart (N=3). Antibodies used: VDAC for loading control; NDUFB8, SDHB, UBCRC2, MtCO1, and ATP5A for ETC complexes I, II, III, IV, and V, respectively.

(J) Confocal imaging showed loss of YAP1 nuclear localization in Group II iPSC-CM.

(K, L) Quantitative analysis of confocal imaging analysis showed no change in total activated β-catenin (K) and YAP1 (L) in iPSC-CM in Group I or II iPSC-CM

Bar graphs show mean ± SEM with one-way ANOVA applied. Box plots show median and minimum-maximum, with Kruskal-Wallis test applied. (B) n=3,6,4 subjects. (C-F) n=3,5,3subjects. (K) n=44,15,17 CMs. (L) n=90,78,79 CMs.

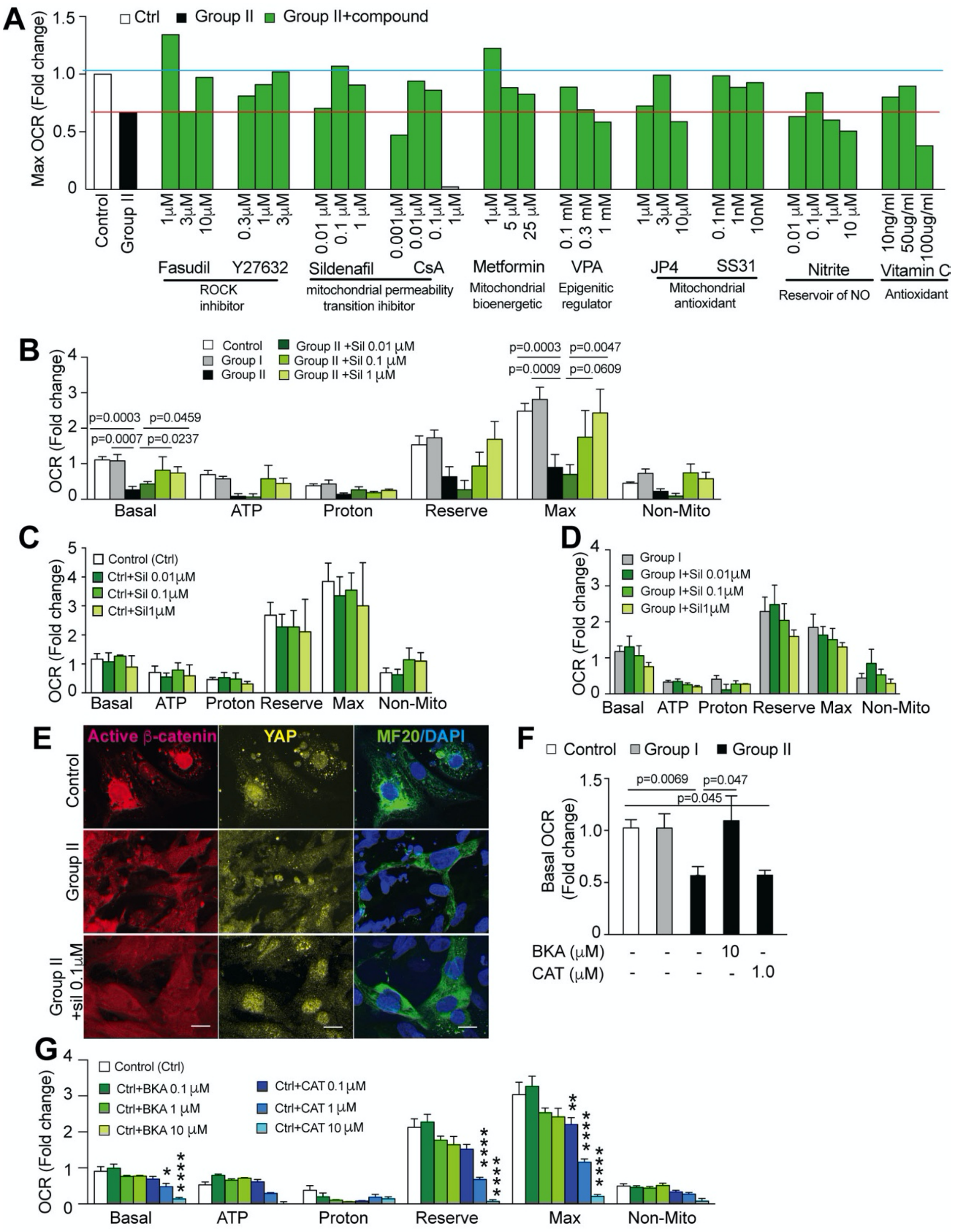


**Figure S4. Inhibition of the mitochondrial permeability transition pore rescues mitochondrial respiration and YAP1 nuclear localization.** Related to Figure 4.

(A) Screening of compounds for potential rescue of the mPTP closure defect was conducted using Group II iPSC-CM with measurement of OCR using the Seahorse Analyzer. Different drug doses were tested to examine efficacy in restoring maximal OCR as compared to the maximal OCR in untreated control and Group I iPSC-CM.

(B-D) Impact of sildenafil on OCR in Group II HLHS iPSC-CM was assessed using the Seahorse Analyzer. Sildenafil rescues mitochondrial respiration in Group II iPSC-CM (B) in a concentration-dependent manner, but has no significant effect on Group I (C) or control (D) iPSC-CM.

(E) Sildenafil treatment of Group II iPSC-CM rescues Yap1 nuclear localization, but not β-catenin, which remained cytoplasmic. (Scale bar =10 mm)

(F) Basal OCR was rescued by bongkrekic acid (BKA), but not not carboxyatractyloside (CAT) in the iPSC-CM from group II HLHS patients.

(G) Control subject iPSC-CM treated with BKA showed no change in mitochondrial respiration, but CAT treatment decreased multiple parameters of mitochondrial respiration.

Bar graphs show mean±SEM, analyzed by one-way ANOVA, n> =3 independent repeats for each bar.

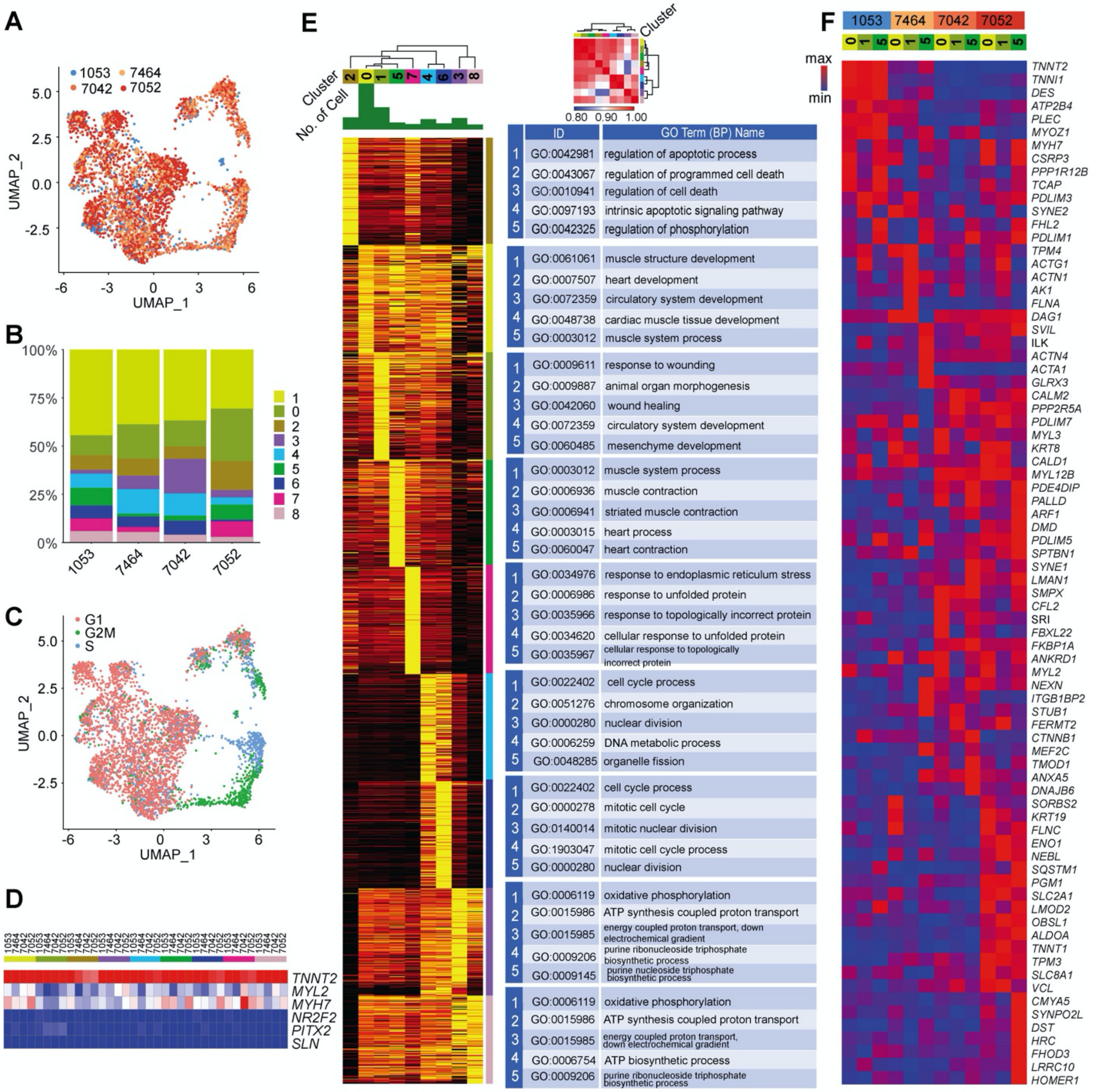


**Figure S5. Analysis of HLHS patient iPSC-CM using single-cell RNAseq** Related to Figure 5.

(A) UMAP plot of single cell RNAseq data obtained for the control subject (1053) and three HLHS patients – Group I patient 7464, and Group II patients 7042 and 7052.

(B) Bar plot showed the proportion of cells in each cluster derived from the three patients and one control subject.

(C) UMAP showing cell cycle distribution among the iPSC-CM.

(D) Heatmap of normalized expression levels of ventricular-like (TNNT2/MYL2/MYH7) and atrial-like marker (NR2F2/PITX2/SLN) showed the iPSC-CM from the control and three HLHS patients are all largely ventricle-type.

(E) Heatmap of normalized expression levels of 200 most significantly upregulated genes in all 8 clusters. Right side denote top 5 GO terms from functional association analysis for each subject across all clusters. On the top right corner, similarity between these clusters were evaluated using Pearson correlation of genes in gene modules. Hierarchical clustering was applied for both rows and columns.

(F) Expression of myofibrillary related genes in Clusters 0,1 and 5 are shown for each sample.

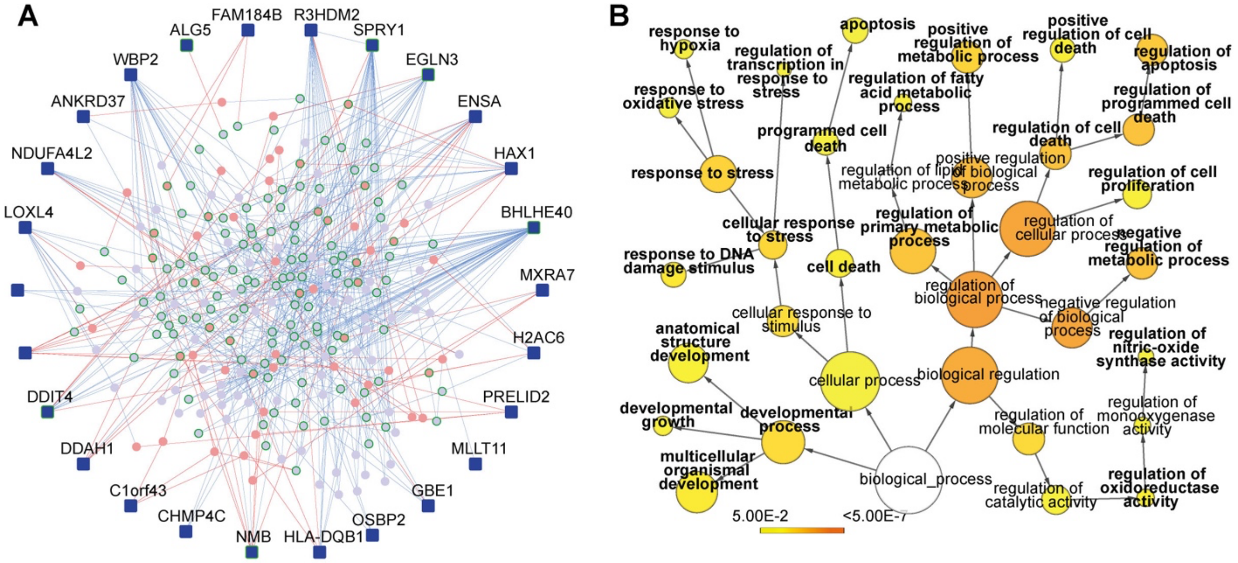


**Figure S6. Protein-Protein Interactome of Genes Highly Expressed Only in Patient 7052.** Related to Figure 5.

(A) There are 28 genes highly expressed only in Patient 7052 shown in region denoted by asterisk in the heatmap of **Figure 5G**. Of these 28 genes, 27 are protein coding genes. These were used to assemble a protein-protein interactome (PPI) network. This yielded a PPI that incorporated 26 of 27 protein coding genes. These 26 genes are shown as nodes (dark blue square) and the protein-protein interactions (PPIs) as edges. Red nodes/edges are novel interactors/interactions. Light blue nodes/edges are known interactors/interactions. Nodes with green-colored borders are genes belonging to selected relevant GO Biological Processes shown in Figure 5I with bold labels.

(B) Protein-protein interactome constructed from the 28 DEGs yielded GO term enrichment visualized using BiNGO, Terms relevant to this work are bolded and their ontological structure from the root are shown. Circle size is proportional to the number of genes in the GO term, and p values are indicated by color, ranging from Yellow (-logP=1.3) to orange (-logP=6.3).

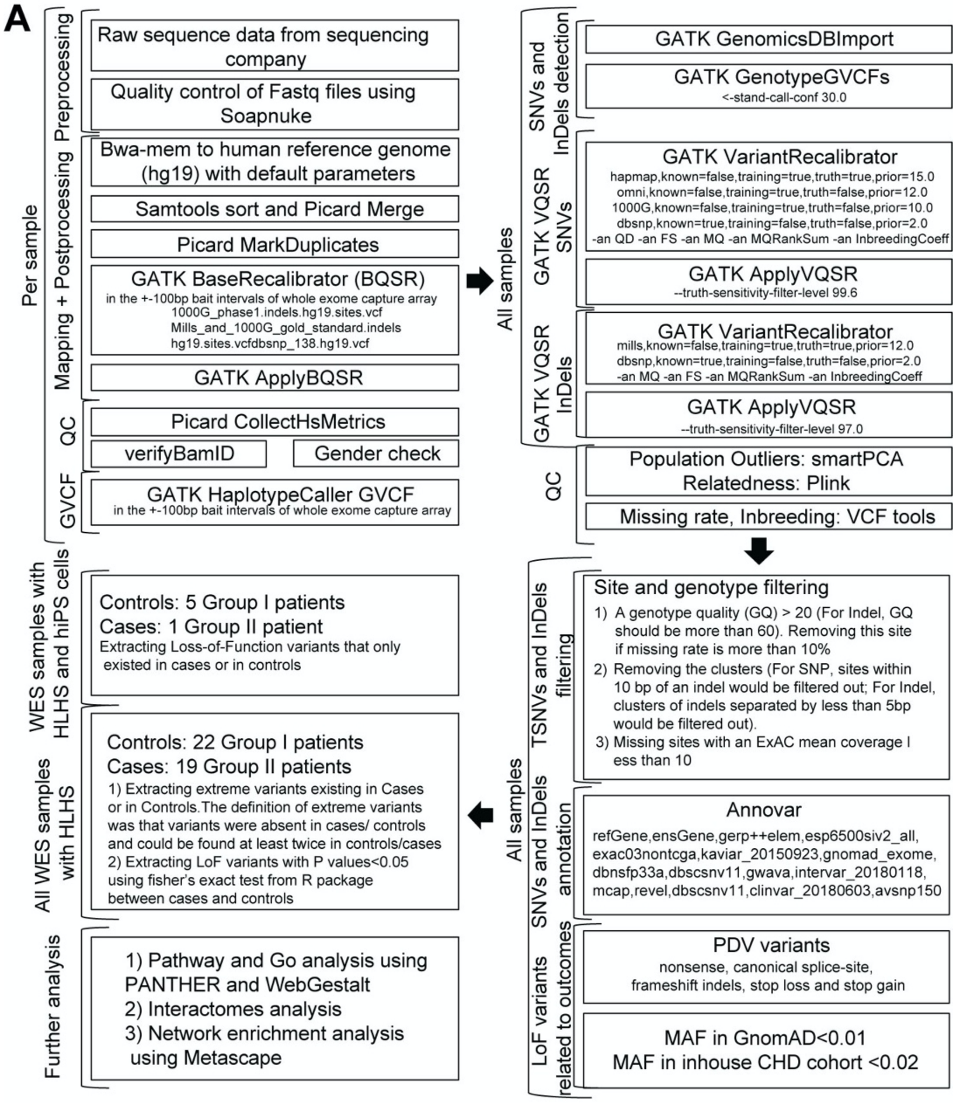


**FigureS7. Whole exome sequencing analysis of pathogenic variants show enrichment for metabolic-mitochondrial pathways in Group II HLHS patients** Related to Figure 7.

**(A)** Bioinformatic workflow for variant detection and filtering.

MAF: minor allele frequency. LoF: Loss of function. DAVID: Database for Annotation Visualization and Integrated Discovery, https://david.ncifcrf.gov/. Metascape: http://metascape.org/gp/index.html#/main/step1. WebGestalt: WEB-based Gene SeT AnaLysis

Toolkit, http://www.webgestalt.org/. ExAC: http://exac.broadinstitute.org/. Annovar: http://annovar.openbioinformatics.org/en/latest/. GnomAD: https://gnomad.broadinstitute.org/.

**SUPPLEMENTAL SPREADSHEET INFORMATION**

**Supplemental Spreadsheet 1: Patient information, iPSC production, antibodies, and primer sequences. Related to STAR Methods.**

14. Toppgene analysis of those 19 overlapping genes (Figure 7D)

### SUPPLEMENTAL VIDEO LEGEND

#### Supplemental Videos

Sup-video-1_hips-cm_beating-Related to Figure1

Sup-video-2_hips-cm_Ca-Related to Figure1

Sup-video-3_hips-cm_single_cell-Related to Figure1

**Sup-video-1_hips-cm_beating:** Videomicroscopy showing contraction of human iPSC-CM. The iPSC-CM from control subject and Group I beat faster than iPSC-CM from Group II. Scale bar = 250 μm.

**Sup-video-2_hips-cm_Ca:** Calcium transients in the iPSC-CM are visualized using Rhod-2. Note faster propagation of calcium transients in iPSC-CM from control subject and Group I patients as compared to that of Group II. Scale bar = 250 μm.

**Sup-video-3_hips-cm_single_cell:** Videomicroscopy recording of individual beating iPSC-CM from control subject, Group I and Group II patients. Robust contractions are seen in cardiomyocytes from control and Group I, but only weak contractions are seen in Group II. Scale bar = 10 μm.
